# Afferent-specific modulation of excitatory synaptic transmission by acetylcholine and serotonin in the prelimbic cortex

**DOI:** 10.1101/2025.05.04.652144

**Authors:** Arielle L. Baker, Allan T. Gulledge

**Affiliations:** Department of Molecular and Systems Biology, Geisel School of Medicine at Dartmouth College, 74 College Street, Vail 601, Hanover, New Hampshire 03755, USA

**Author notes:** Author contributions: A.L.B. contributed to experimental design, data acquisition, analysis, interpretation of data, and edited the manuscript. A.T.G. conceived of the project and contributed to experimental design, data acquisition, analysis, interpretation of data, and wrote and edited the manuscript. The authors declare no competing financial interests.

## Abstract

Acetylcholine (ACh) and serotonin (5-hydroxytryptamine, or 5-HT) differentially regulate the excitability of pyramidal neurons in the mouse prelimbic (PL) cortex according to their long-distance projections. Here we tested for afferent- and/or target-specific modulation of glutamate release by ACh and 5-HT in two long-distance excitatory projections to the PL cortex: commissural (COM) afferents from the contralateral cortex and projections from the mediodorsal nucleus (MDN) of the thalamus. Using ex-vivo optogenetic approaches, we mapped the connectivity and neuromodulation of COM and MDN afferents in layer 5 intratelencephalic (IT) and extratelencephalic (ET) projection neurons. Dual whole-cell recordings in pairs of IT and ET neurons revealed that COM afferents target both neuron subtypes, but that MDN afferents selectively target IT neurons. Both afferents exhibited similar target-independent short-term synaptic plasticity (paired-pulse facilitation) across a range of frequencies, but were differentially modulated by ACh and 5-HT. In both control conditions and after isolating monosynaptic connections with tetrodotoxin and 4-aminopyridine, COM transmission was suppressed strongly by ACh and moderately by 5-HT, while MDN transmission was largely unaffected by either neuromodulator. Suppression o,f COM transmission by ACh or 5-HT was concentration dependent and mediated by M4 muscarinic or 5-HT1B receptors, respectively. Chemogenetic inhibition of hM4Di-expressing COM terminals mimicked the suppressive effects of ACh and 5-HT on synaptic transmission. Our results demonstrate that ACh and 5-HT preferentially regulate COM synaptic transmission, albeit to different degrees, and suggest that, through their combined pre- and postsynaptic neuromodulation, ACh and 5-HT may differentially regulate cortico-striatal-thalamic loops to influence cognition and behavior.

## Introduction

In the mouse prelimbic (PL) cortex, acetylcholine (ACh) and serotonin (5-HT) reciprocally modulate the excitability of two layer 5 pyramidal neuron subtypes: intratelencephalic (IT) neurons that project bilaterally within the cerebrum, and extratelencephalic (ET) neurons that project unilaterally to the midbrain and brain-stem (Tudi et al., 2024). Whereas ACh preferentially enhances the excitability of ET neurons via Gq-coupled M1-type muscarinic ACh receptors (mAChRs; Gulledge et al., 2009; Dembrow et al., 2010; Baker et al., 2018), 5-HT acts at Gi/o-coupled 5-HT1A (1A) receptors to inhibit ET neurons while simultaneously increasing IT neuron excitability via Gq-coupled 5-HT2A (2A) receptors (Avesar and Gulledge, 2012; Stephens et al., 2014; Stephens et al., 2018). Thus, cortical release of ACh is expected to enhance ET-dependent corticofugal output, whereas 5-HT release may inhibit corticofugal output while amplifying corticocortical and corticostriatal circuits. However, because presynaptic ACh and 5-HT receptors may also regulate the excitatory synaptic transmission that drives pyramidal neurons (e.g., Murakoshi et al., 2001; for reviews, see Schicker et al., 2008; Betke et al., 2012; Yang et al., 2021), predicting the net effect if a given neuromodulator on cortical circuit function will require a detailed understanding of both pre- and postsynaptic mechanisms, and their interactions, across cortical neuron subtypes and local and long-distance afferents.

Two important excitatory afferents to the PL cortex include commissural (COM) afferents from the contralateral PL cortex and inputs from the mediodorsal nucleus of the thalamus (MDN). To date, few studies have examined the effects of ACh or 5-HT on optogenetically isolated excitatory afferents to the PL cortex, and none have compared modulation of COM or MDN synaptic transmission across IT and ET target neurons. Only one previous study has tested for serotonergic regulation of glutamate release in an optogenetically isolated afferent in the PL cortex, reporting that 5-HT moderately suppressed COM transmission in the mouse PL cortex (Kjaerby et al., 2016). Similarly, only one prior study has tested for cholinergic modulation of an optogenetically isolated afferent, finding that inputs from the nucleus reuniens to the PL cortex were insensitive to ACh (Banks et al., 2021). However, neither of these studies tested for target-neuron-specific modulation of monosynaptic excitatory transmission.

In the mouse anterior cingulate cortex, excitatory postsynaptic currents (EPSCs) evoked by electrical stimulation of putative COM afferents were modestly suppressed by 5-HT (Troca-Marin and Geijo-Barrientos, 2010; see also Tian et al., 2017). On the other hand, ACh was found to bidirectionally modulate synaptic transmission in superficial pyramidal neurons in the PL cortex, with mAChRs suppressing, but nicotinic-type ACh receptors enhancing, electrically evoked excitatory postsynaptic potentials (EPSPs; Vidal and Changeux, 1993). Muscarinic and/or serotonergic suppression of synaptic transmission has also been reported for electrically evoked (Gil et al., 1997; Hsieh et al., 2000; Atzori et al., 2005) and unitary (Tsodyks and Markram, 1997; Levy et al., 2006, 2008; Yang et al., 2020; Agahari and Stricker, 2021, 2025) connections in the sensory cortexes of young (typically < 1 month-old) rodents.

Less is known regarding neuromodulation of thalamic inputs to the PL cortex. In layer 5 neurons, 5-HT was reported to enhance electrically evoked, putative MDN synaptic responses via presynaptic 2A receptors (Barre et al., 2016) whereas a lack of cholinergic modulation was reported in afferents from the nucleus reuniens (Banks et al., 2021). In the somatosensory cortexes of neonatal or very young rodents, 5-HT suppressed putative thalamocortical transmission, an effect that was reduced (Rhoades et al., 1994) or eliminated (Laurent et al., 2002) in adolescence, whereas activation of 2A receptors enhanced thalamocortical glutamate release in the adult retrosplenial cortex (Ekins et al., 2025). On the other hand, ACh bidirectionally modulated putative thalamocortical synapses via muscarinic and nicotinic receptors in somatosensory cortexes from juvenile mice and rats (Gil et al., 1997; see also Hsieh et al., 2000).

Given the rich substrate for potential afferentand/ or target-specific presynaptic modulation of excitatory synaptic transmission in the PL cortex, and the reciprocal nature of postsynaptic cholinergic and serotonergic modulation of ET and IT neuron excitability, we asked whether these same neuromodulators may exhibit afferent- and/or target-neuron-specific regulation of excitatory synaptic transmission. Using an optogenetic approach, we compare, for the first time, neuromodulation of glutamate release by ACh and 5-HT in optogenetically triggered and pharmacologically isolated monosynaptic COM and MDN inputs to IT and ET neurons.

## Methods

### Animals and ethical approvals

Experiments utilized female and male 7-to 32-week-old C57BL/6J mice according to protocols approved by the Institutional Animal Care and Use Committee of Dartmouth College. Mice were bred and cared for in dedicated facilities accredited by the Association for Assessment and Accreditation of Laboratory Animal Care. With food and water continuously available, mice experienced a 12-hour light-dark cycle and were housed with littermates until surgery (see below). Post surgery, animals were housed individually until use in experiments.

### Animal surgeries

Intracranial injections were used to deliver adeno-associated viruses (AAVs) encoding channelrhodopsin-2 (ChR2) or chemogenetic tools to specific neuron populations. During surgeries, mice were anesthetized with vaporized isoflurane (~2%), given an intraperatoneal injection of ketoprofen as an analgesic (4 mg/kg), and secured in a stereotaxic frame. Craniotomies and intracerebral injections were made according to appropriate coordinates (**Table 1**), with microsyringes lowered into the brain region of interest (i.e., medial prefrontal cortex [mPFC] or mediodorsal nucleus [MDN] of the thalamus), and infusions of AAV viruses made at a rate of 50 nL per minute. Mice were allowed to recover from surgery for at least 3 weeks prior to use in electrophysiology experiments. Accuracy of injection sites was confirmed via visualization of mCherry in coronal brain sections of the PL cortex or thalamus.

**Table l.** AAV manipulations of prelimbic afferents.

| Target afferent | Hemisphere (relative to recording site) | Coordinates (mm; relative to bregma) | Addgene # | Virus | Dilution | volume (nL) |
| --- | --- | --- | --- | --- | --- | --- |
| Commissural (COM) | contralateral | lateral: ±0.48<br>rostral-caudal: +2.10<br>depth: -1.60 | 100054 | pAAV.CAG.hChR2(H134R)-mCherry.WPRE.SV40 | 1:10 (*1:5) | 300 (*150) |
|  |  |  | 50475 | pAAV-hSyn-hM4D(Gi)-mCherry | 1:5 | 250 |
|  |  |  | 50474 | pAAV-hSyn-hM3D(Gq)-mCherry | 1:5 | 250 |
| Mediodorsal nucleus (MDN) | ipsilateral | lateral: ±0.70<br>rostral-caudal: -1.48<br>depth: -3.30 | 100054 | pAAV.CAG.hChR2(H134R)-mCherry.WPRE.SV40 | 1:10 | 300 |

All AAVs were of the AAV9 serotype. \*When ChR2 was co-expressed with hM4Di or hM3Dq in COM afferents, dilution and concentration were reduced by half.

### Slice preparation

Anesthetized mice (2% vaporized isoflurane) were decapitated and their brains removed rapidly into carbogen-(95% O2 / 5% CO2) saturated artificial cerebral spinal fluid (aCSF) composed of (in mM): 125 NaCl, 25 NaHCO3, 3 KCl, 1.25 NaH2PO4, 0.5 CaCl2, 6 Mg-Cl2, and 25 glucose. Coronal slices (250 µm thick) were cut with a vibratome (Leica VT 1200) and stored in holding chambers filled with aCSF containing 2 mM CaCl2 and 1 mM MgCl2. Slices were left to recover for ~45 minutes at 35°C and then stored at room temperature (~26°C) until use in experiments.

### Electrophysiology

Slices were transferred to a low-volume (~1 ml) recording chamber perfused continuously (~5 ml / min) with warm (35 - 36°C) carbogen-saturated aCSF under an Olympus upright microscope. Using a 60x objective and oblique illumination, somata of layer 5 cortical neurons in the PL cortex (330 - 500 µm from the pia) were visually targeted for whole-cell current-clamp recording. Patch pipettes (5 - 7 MΩ) were filled with a solution containing (in mM): 135 potassium gluconate, 2 NaCl, 2 MgCl2, 10 HEPES, 3 Na2ATP, and 0.3 NaGTP, pH 7.2 with KOH. Recordings were achieved with either BVC-700 amplifiers (Dagan Corporation) and Axograph software (AxoGraph Company) or dPatch amplifiers running SutterPatch software in Igor Pro (Sutter Instrument). Electrode capacitance and series resistance were maximally compensated and membrane potential sampled at a minimum of 25 kHz (filtered at 5 or 10 kHz) and corrected for the liquid junction potential of +12 mV. Use of current-clamp allowed the physiological classification of IT and ET neurons, action potential generation, and avoided the significant distortion of synaptic events that occurs when voltage-clamp is attempted in pyramidal neurons (e.g., Williams and Mitchell, 2008; Dembrow et al., 2015; Beaulieu-Laroche and Harnett, 2018).

To control for variable levels of ChR2 expression across mice, and to determine relative strength of excitatory afferents to specific postsynaptic neuron sub-types, in many experiments simultaneous dual recordings were made of IT and ET neurons (typically within 75 µm of each other). IT and ET neurons were identified as previously described according to their physiological characteristics (Gulledge, 2024; see also Elliott et al., 2018). Briefly, neurons were classified based on a “Physiology Index” (PI) determined from physiological properties that are influenced by the differential expression of hyperpolarization-activated and cyclic nucleotide-gated (HCN) channels in IT and ET neurons (Dembrow et al., 2010; e.g., Sheets et al., 2011; Avesar and Gulledge, 2012; Oswald et al., 2013). The PI was computed for each neuron according to the following formula:

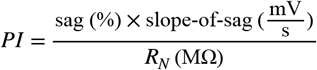

where RN is the neuron’s input resistance (in MΩ), “sag” is the relative magnitude of the steady-state HCN-channel-mediated rebound in membrane potential during a ~20 mV (peak) hyperpolarizing current injection (as a percent of the peak hyperpolarization), and “slope-of-sag” is the rate of voltage rebound (mV/s) as determined by linear regression over a 50 ms period beginning at the time of peak hyperpolarization. Neurons with a PI of greater than 4.5 were identified as ET, while those with a PI lower than 3.5 were identified as IT (**Table 2**). Neurons with intermediate PIs were excluded from analysis (Gulledge, 2024).

**Table 2.** Physiological classification of IT and ET neurons in prelimbic cortex.

| Neuron subtype | n | Age (days) | $V_M$ (mV) | $R_N$ (MΩ) | Sag (%) | Slope of sag (mV/s) | Physiology index |
| --- | --- | --- | --- | --- | --- | --- | --- |
| IT | 287 | 92 ± 37 | -83.2 ± 4.9 | 127 ± 50 | 7.5 ± 4.6 | 9.9 ± 9.9 | 0.8 ± 1.4 |
| ET | 308 | 90 ± 34 | -78.0 ± 3.2 | 52 ± 17 | 19.6 ± 6.4 | 49.7 ± 17.5 | 21.7 ± 15.3 |
| IT vs ET | p-value | 0.425 | <0.001 |  |  |  |  |
|  | Cohen’s *d* | 0.07 | 1.27 |  |  |  |  |
| IT female | 111 | 95 ± 35 | -83.2 ± 5.1 | 123 ± 49 | 7.7 ± 4.9 | 9.5 ± 8.5 | 0.9 ± 1.9 |
| IT male | 176 | 90 ± 38 | -83.2 ± 4.8 | 130 ± 51 | 7.4 ± 4.3 | 10.1 ± 10.7 | 0.8 ± 0.9 |
| IT female vs IT male | p-value | 0.213 | 0.950 | 0.243 | 0.607 | 0.612 | 0.592 |
|  | Cohen’s *d* | 0.15 | 0.01 | 0.14 | 0.06 | 0.06 | 0.08 |
| ET female | 139 | 91 ± 33 | -78.0 ± 3.3 | 51 ± 18 | 19.2 ± 6.1 | 48.9 ± 15.6 | 21.3 ± 15.6 |
| ET male | 169 | 85 ± 32 | -78.0 ± 3.0 | 52 ± 16 | 19.9 ± 6.6 | 50.4 ± 18.9 | 22.0 ± 15.1 |
| ET female vs ET male | p-value | 0.012 | 0.873 | 0.414 | 0.345 | 0.435 | 0.691 |
|  | Cohen’s *d* | 0.30 | 0.02 | 0.09 | 0.11 | 0.09 | 0.05 |

Physiological properties of female and male neurons classified as IT or ET shown as means ± standard deviations. $V_M$ is the resting membrane potential, $R_N$ is the input resistance, sag and slope of sag are measures of HCN-channel mediated rebound from peak hyperpolarizations (see Methods). *P*-values are for Student’s *t*-tests and effect sizes are represented by Cohen’s *d*. Statistical comparisons were not made across neuron subtypes for metrics used to define those subtypes (see Methods).

EPSPs were optically evoked using LED-driven epifluorescent flashes of blue light (460 or 470 nm; 1 ms) with the 60x objective over layer 5. LED intensities were measured at the objective with a LaserCheck intensity meter (model 45-018; Edmond Optics). EPSPs were typically evoked in pairs at 20 Hz, with trials repeated at 15 s intervals to allow for full recovery from short-term plasticity. Measurements of EPSP characteristics, including amplitudes and short-term plasticity (e.g., paired-pulse ratios; PPRs), were made from averaged responses of ten consecutive trials in each experimental condition. EPSP amplitudes and PPRs were measured relative to the mean resting membrane potential in the 50 ms prior to the initial light flash, while measurements of EPSP onset latency (determined at 5% amplitude) were made relative to the onset of light flashes. EPSP width (which is primarily a measure of decay kinetics) was made at half of the peak amplitude relative to the baseline membrane potential.

In most experiments, LED power was adjusted such that the first of the two EPSPs was ~10 mV in amplitude while still allowing for a subthreshold second EPSP. For comparisons of afferent targeting, only IT-ET pairs in which at least one neuron had an EPSP ≥ 5 mV were used for analyses. Baseline somatic resting membrane potentials (**Table 2**) were maintained throughout experiments using slow DC current injection (manually, or using the dynamic holding function of the dPatch). Pharmacological agents (e.g., ACh or 5-HT) were bath applied for 7 minutes with measurements made from the average of the final 10 EPSPs in each experimental condition. For each synapse, the coefficient of variation (CV) of EPSP amplitudes was determined from 10 or 12 consecutive EPSPs in each experimental condition. Pharmacological manipulations were followed by a 15 - 20 minute wash period. For dose-response experiments, normalized mean data were fit with a Hill equation using Igor Pro’s CurveFit function constrained to a presumed null effect at 1 pM concentration. For data with ACh, where all concentrations except for “eserine-only” generated suppression of EPSPs greater than 50%, we first fit the raw data with an initial Hill equation and then used that curve to estimate the eserine-only concentration of ACh (139 nM), before fitting a final Hill equation to the mean data including the eser-ine-only estimate.

### Pharmacological agents

Most substances were dissolved in water to make stock solutions (typically at 10,000x concentration) that were frozen as aliquots and used as needed during experiments. Where necessary, stock solutions were made with DMSO at maximal solubility. Acetylcholine (Fisher Scientific) was bath-applied with 10 µM physostigmine hemisulfate (eserine; Tocris Bioscience), a blocker of acetylcholinesterase. Atropine, a nonspecific muscarinic antagonist, was purchased from Sigma Aldrich. Antagonists for M4 (VU 6028418 hydrochloride) and M2 (AF-DX 116) receptors were purchased from Tocris Biosciences, while the M4 positive allosteric modulator VU0467154 was purchased from MedChemExpress. Antagonists for 5-HT receptors included SB-216641 (a 5-HT1B receptor antagonist; Abcam), SB-224289 (a second 5-HT1B receptor antagonist; Tocris Bioscience), and SB-699551 (a 5-HT5A receptor antagonist; Santa Cruz Biotechnology). 5-HT1 receptor agonists included CP-93129 (targeting 1B receptors; Tocris Bioscience), PNU-142633 (targeting 5-HT1D receptors; Tocris Bioscience), and BRL-54443 (targeting 5-HT1F receptors; MedChemExpress). Tetrodotoxin (TTX) citrate and clozapine N-oxide (CNO) hydrochloride were purchased from HelloBio, Inc., whereas 4-aminopyridine (4-AP) was purchased from Fisher Scientific.

### Statistical analysis

Unless otherwise noted, data are presented as mean values ± standard deviation. Two-tailed Student’s t-tests for paired or unpaired data were used, as appropriate, to compare physiological responses across experimental condition or across neuron subtypes, but no assumptions were made regarding the “significance” of differences in means (see Nieuwenhuis et al., 2011; e.g., Wasserstein et al., 2019). Instead, we quantified effect size as Cohen’s *d*, a measure of standardized mean difference, according to the following formulas:

for non-paired data:

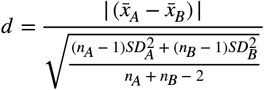

where 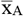 and 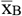 are the mean values of a given parameter (e.g., EPSP amplitude), SDA and SDB are the standard deviations for those parameters, and nA and nB are the number of observations (i.e., neurons) in each group, or, for paired data, the lessor of:

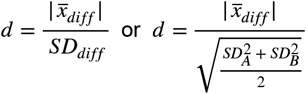

where 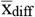 is the mean paired difference of a parameter and SDdiff is the standard deviation of the differences. The two equations differ with respect to the denominator, and to be as conservative as possible, the lessor of these values for each data set is reported. By convention, effect sizes ≥ 0.8 are considered “large” (i.e., having distributions that overlap by ~53% or less, by area), whereas values of ~0.5 and ~0.2 are considered to be “medium” or “small” effects, respectively, according to their greater (~67% and ~85%) overlap of distribution area (Cohen, 1988). Wherever possible, statistics and effect sizes are presented in the appropriate data tables below, rather than in the main text.

## Results

### Targeting of IT and ET neurons by commissural and thalamocortical afferents

Dual whole-cell recordings and ChR2-assisted circuit mapping (Petreanu et al., 2007) was used to quantify the relative targeting of IT and ET neurons by commissural (COM; n = 99 IT-ET pairs) or thalamocortical (MDN; n = 58 IT-ET pairs) afferents to the PL cortex (**Figure 1A-D**; **Table 3**). Afferent-specific synaptic transmission was triggered with two brief (1 ms duration) flashes of blue light with a 50 ms inter-flash interval (20 Hz), repeated every 15 s. Flash-evoked EPSPs were sensitive to LED intensity (**Figure 1C, E**) and, with the exception of MDN inputs to ET neurons, had short (< 2 ms) onset latencies (**Table 3**). When LED intensities were adjusted to generate, as closely as possible, ~10 mV initial EPSPs across both neurons, while still allowing for subthreshold EPSPs on the second flash, COM EPSPs in IT neurons were slightly larger than those recorded simultaneously in ET neurons, while PPRs for COM inputs were similar across target neuron subtypes (**Figure 1D, E**; **Table 3**). These data suggest that COM afferents robustly form synapses with similar short-term plasticity among layer 5 IT and ET target neurons in the PL cortex.

**Table 3.** Properties of afferent inputs to pairs of IT and ET neurons in the prelimbic cortex.

| Afferent<br>(# of<br>pairs) | Target neuron<br>subtype | EPSP 1 |  |  | EPSP 2 |  |  | PPR |
| --- | --- | --- | --- | --- | --- | --- | --- | --- |
|  |  | Amplitude<br>(mV) | Onset<br>latency<br>(ms) | half-<br>width<br>(ms) | Amplitude<br>(mV) | Onset<br>latency<br>(ms) | half-<br>width<br>(ms) |  |
| COM<br>(99) | IT | 10.4 ± 4.2 | 1.88 ± 0.41 | 21 ± 8 | 15.6 ± 6.4 | 1.79 ± 0.41 | 25 ± 11 | 1.52 ± 0.36 |
|  | ET | 9.0 ± 4.0 | 1.89 ± 0.40 | 14 ± 4 | 13.6 ± 5.9 | 1.89 ± 0.37 | 16 ± 5 | 1.56 ± 0.41 |
|  | IT p-value<br>vs ET | 0.019 | 0.685 | < 0.001 | 0.011 | < 0.001 | < 0.001 | 0.335 |
|  | Cohen's *d* | 0.24 | 0.02 | 0.91 | 0.26 | 0.28 | 0.99 | 0.10 |
| MDN<br>(58) | IT | 15.6 ± 8.4 | 1.98 ± 0.54 | 27 ± 10 | 21.3 ± 10.0 | 1.96 ± 0.53 | 28 ± 12 | 1.44 ± 0.41 |
|  | ET | 6.4 ± 3.9 | 3.03 ± 1.33 | 20 ± 7 | 11.4 ± 6.4 | 2.97 ± 1.12 | 23 ± 8 | 1.94 ± 0.86 |
|  | IT p-value<br>vs ET | < 0.001 | < 0.001 | < 0.001 | < 0.001 | < 0.001 | < 0.001 | < 0.001 |
|  | Cohen's *d* | 0.94 | 0.83 | 0.69 | 0.86 | 0.82 | 0.49 | 0.63 |

Mean EPSP amplitudes (in mV) and paired-pulse ratios (PPR; ± standard deviations) for optically triggered commissural (COM) and thalamocortical (MDN) synaptic events recorded in pairs of IT and ET target neurons. *P*-values are for Student's *t*-tests for paired samples. Effect sizes are represented by Cohen's *d*.

**Figure 1.**
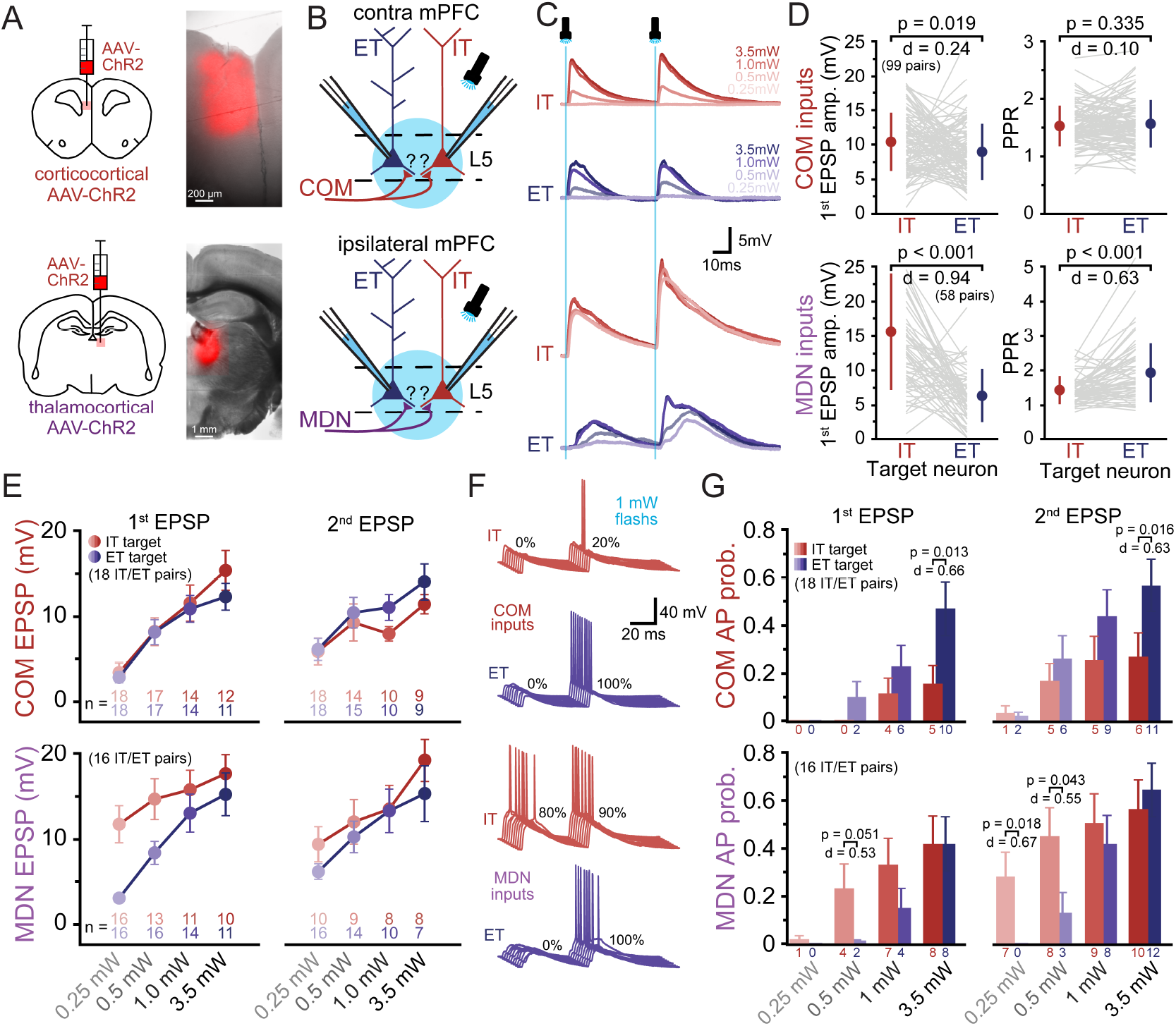
Corticocortical and thalamocortical afferent input to IT and ET neurons in the prelimbic cortex. ***A***, Diagram (left) of approach for delivering ChR2-mCherry to commissural (COM) corticocortical (top) and thalamocortical (MDN; bottom) afferents to the prelimbic cortex, and images of injection sites (right). ***B***, Diagrams of experimental setup for recording light-triggered COM (top) or MDN (bottom) EPSPs in pairs of IT and ET neurons. ***C***, COM (top) and MDN (bottom) EPSPs (averages of 10 consecutive trials) recorded in pairs of IT (red traces) and ET (blue traces) layer 5 neurons in response to two flashes of blue light (1 ms each, delivered at 20 Hz). ***D***, Comparison of the first EPSP amplitudes (left) and paired-pulse ratios (PPR; right) in pairs of IT (red) and ET (blue) target neurons. Mean values and standard deviations are offset with filled symbols. ***E***, Comparison of COM (top) and MDN (bottom) mean EPSP amplitudes in response to increasing LED power. The number of data points (neurons) falls off at higher light intensities as some neurons had only suprathreshold EPSPs. ***F***, Ten consecutive trials of COM (top) or MDN (bottom) synaptic activation (1 mW) recorded in pairs of IT (red) and ET (blue) neurons. The proportions (%) of trials exhibiting an AP for each flash are shown. ***G***, Plots of the mean probabilities of action potential (AP) generation (± SEM) across all trials and all IT and ET target neurons in response to increasing LED intensity for COM (top) and MDN (bottom) inputs. The number below each bar indicates the number of IT (red) or ET (blue) neurons (out of the 18 or 16 neurons per group) that exhibited an AP in at least one trial at the given LED intensity.

Selective activation of ChR2-expressing MDN afferents generated target-specific responses in IT-ET neuron pairs (**Figure 1C, D**). Flash-evoked MDN EPSPs had faster onset latencies in IT neurons (~2 ms) relative to ET neurons (~3 ms; **Table 3**) that were similar to those observed for COM afferents in both IT (n = 157; p = 0.219, *d* = 0.22) and ET (p = 0.257, *d* = 0.20) target neurons. Onset latencies for MDN EPSP in ET neurons were delayed relative to those in IT neurons (**Table 3**) or relative to COM EPSPs in either IT (p < 0.001; *d* = 1.32) or ET (p < 0.001, *d* = 1.31) target neurons. MDN EPSPs measured in ET neurons were also smaller in amplitude and exhibited greater paired-pulse facilitation than MDN EPSPs occurring in IT neurons (**Figure 1D, E**; **Table 3**) or COM EPSPs recorded in IT (p < 0.001; *d* = 0.70) or ET (p < 0.001; *d* = 0.63) neurons.

The longer onset latencies and greater PPRs for MDN EPSPs ET neurons suggest that MDN-evoked responses in ET neurons may result from polysynaptic excitatory drive within the local cortical network (see also Collins et al., 2018). Indeed, flash-evoked EPSPs were often sufficient to drive action potentials (APs) in postsynaptic neurons, especially at higher LED intensities or following the second of two flashes (**Figure 1F, G**). In 18 IT-ET neuron pairs experiencing flash-evoked COM inputs across a range of LED intensities, APs were rarely observed at lower intensities but became progressively more likely as intensities were increased. At the highest LED intensity, APs were more likely to occur in ET neurons (**Figure 1G**). On the other hand, activation of MDN inputs was more likely to generate APs in IT neurons at low LED intensities, but was equally potent in driving APs in IT and ET neurons at the highest LED intensity (n = 16 IT-ET pairs; **Figure 1G**).

The results described above suggest that optical activation of either COM or MDN afferents may trigger local network activity that contributes to flash-evoked EPSPs. To isolate monosynaptic targeting of IT and ET neurons by COM and MDN afferents, we used TTX (1 µM) to block network activity and added 4-AP (100 µM; Petreanu et al., 2009) and increased LED intensities to allow direct ChR2-evoked release from synaptic terminals (**Figure 2A**; see also below). In the presence of TTX and 4-AP, monosynaptic COM EPSPs were robust in both IT and ET target neurons, being slightly larger in the former (**Figure 2B**), whereas PPRs in both target neuron subtypes were substantially reduced relative to those observed in baseline conditions (**Figure 2C**; **Table 4**). The drop in PPRs in the presence of TTX and 4-AP, which was also reported by Cruikshank et al. (2010), likely results from the slow kinetics of direct ChR2-mediated depolarization of synaptic terminals. Monosynaptic COM EPSPs evoked in the presence of TTX and 4-AP exhibited more variable EPSP amplitudes, as reflected in larger (+130 ± 130%) CVs (from 0.14 ± 0.06 to 0.29 ± 0.17; n = 22 IT and ET neurons; p < 0.001, *d* = 0.91) of EPSP amplitudes after the addition of TTX and 4-AP.

**Table 4.**
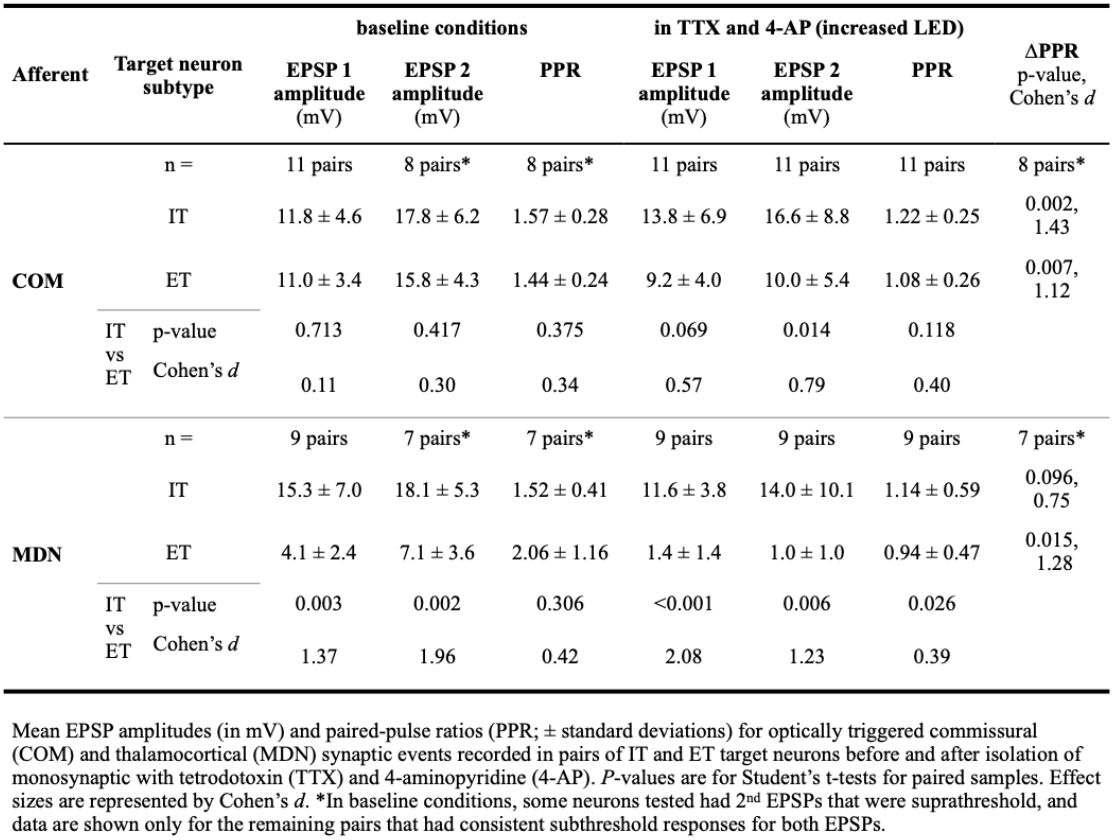
Monosynaptic targeting of IT and ET neurons.

**Figure 2.**
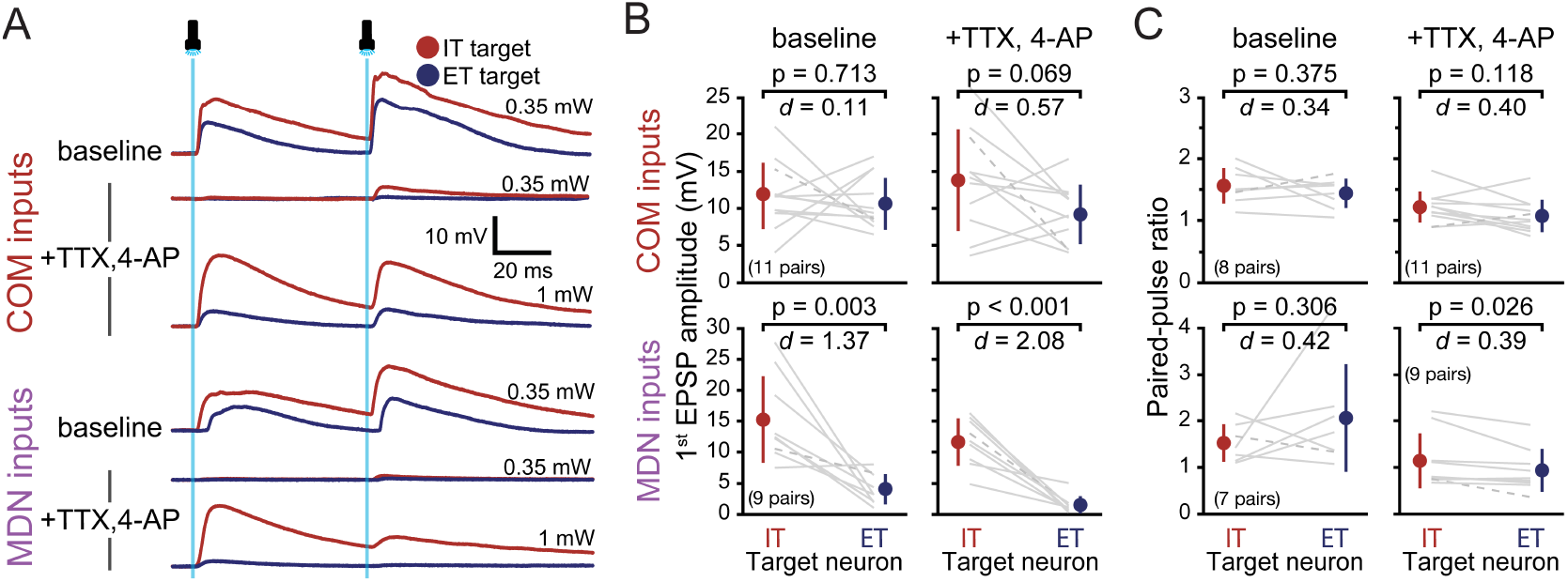
Monosynaptic connectivity of commissural (COM) and thalamocortical (MDN) inputs to IT and ET neurons. ***A***, Average EPSPs (10 consecutive trials) in pairs of IT (red traces) and ET (blue traces) target neurons following activation of COM (upper traces) or MDN (lower traces) afferents in baseline conditions and after application of 1 µM TTX and 100 µM 4-AP at the indicated LED intensities. ***B***, Comparisons of first EPSP amplitudes in IT-ET pairs in baseline conditions and after isolation of monosynaptic inputs with TTX and 4-AP. Gray lines indicate individual neuron pairs, with the dashed lines indicating data for neurons shown in *A*. Mean values and standard deviations are offset with filled symbols. P-values for Student’s t-tests for paired data and effect sizes (Cohen’s *d*) are shown. ***C***, Similar to ***B***, but for paired-pulse ratios. In some neurons, the second EPSP was suprathreshold in baseline conditions, leading to a reduction in the number of pairs quantified for PPR in baseline conditions.

When we isolated monosynaptic MDN inputs with TTX and 4-AP, light-evoked responses in ET neurons became very small and were sometimes undetectable, while EPSPs remained substantial in IT neurons (**Figure 2A, B**; **Table 4**). As was true for COM inputs, the addition of TTX and 4-AP reduced PPRs for MDN EPSPs (**Figure 2C**; **Table 4)** and increased the CV of EPSP amplitudes (by +143 ± 179%, from 0.17 ± 0.16 to 0.31 ± 0.34; n = 18 IT and ET neurons; p = 0.002, *d* = 0.50). These data confirm that MDN-evoked responses in ET neurons are largely polysynaptic in the absence of TTX.

### Frequency dependence of short-term plasticity in COM and MDN afferents

To explore the frequency dependence of short-term synaptic plasticity, in additional neurons (not necessarily dual recordings) we optically evoked trains of ten COM (IT and ET targets) or MDN (IT targets only) EPSPs at 10, 20, or 40 Hz (**Figure 3A, B**). For both COM and MDN inputs, and across all frequencies and target neuron subtypes, initial responses exhibited facilitation of similar magnitude peaking during the 2nd (for 10 Hz), 2nd or 3rd (for 20 Hz), or 3rd - 5th (for 40 Hz) light flashes, before becoming smaller toward the end of the trains. These results are comparable to those of Martinetti et al., (2022), who observed paired-pulse facilitation of long-distance ipsilateral corticocortical connections that peaked at ~20 Hz.

**Figure 3.**
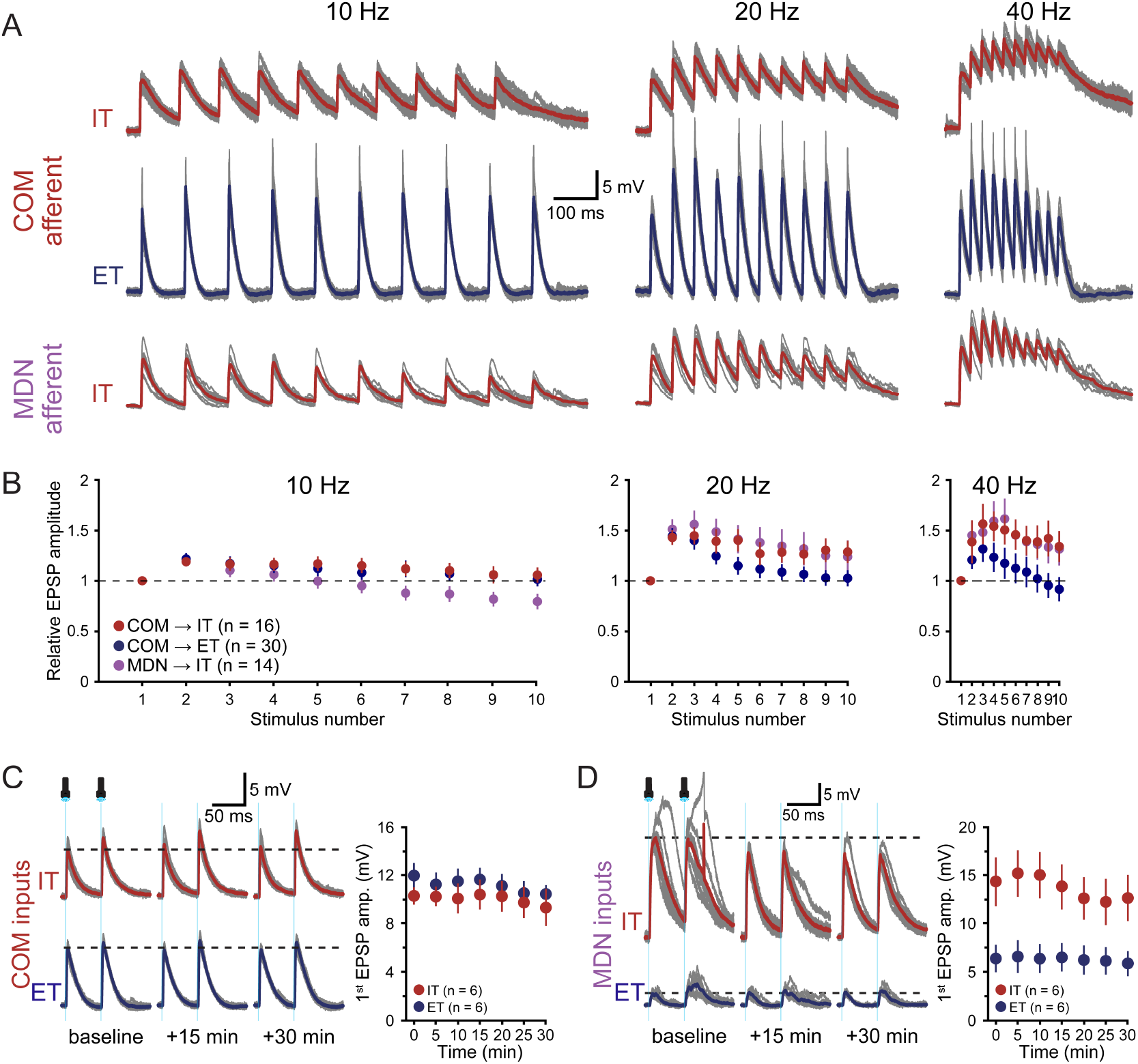
Frequency-dependent short-term plasticity in commissural (COM) and corticothalamic (MDN) inputs to IT and ET neurons and response stability over time. ***A***, Ten consecutive responses (gray traces) and their averages (colored traces) to trains of 10 flashes delivered at 10 (left), 20 (middle), and 40 Hz (right) Hz in a IT-ET pair of target neurons experiencing COM input (top and middle traces) and in a different IT neuron experiencing activation of MDN inputs (lower traces). ***B***, Plots of mean normalized EPSP amplitudes (relative to 1^st^ EPSP; ±SEM) during trains of ten optically evoked EPSPs from COM afferents onto IT and ET neurons (red and blue symbols, respectively; not all data from pairs) and from MDN afferents in IT neurons (purple symbols) at the indicated frequencies. ***C***, Ten consecutive trials (gray traces) and average responses (colored traces) in an IT-ET pair experiencing COM-evoked synaptic input under baseline conditions over a 30 minute period (left) and a summary plot of mean EPSP amplitudes over time (±SEM) for 6 IT and 6 ET neurons (not necessarily in pairs). ***D***, Similar to ***C***, but for optical activation of MDN inputs.

At 10 Hz, we observed in IT neurons a difference in the short-term plasticity of COM and MDN inputs during the second half of the train, with MDN inputs generating relatively smaller late EPSPs relative to the initial EPSP (p ranging from 0.065 to 0.022 and *d* ranging from 0.78 to 0.88 over the final five EPSPs; **Figure 3B**). This afferent-specific difference in short term plasticity was not apparent in trains generated at 20 or 40 Hz in the same neurons. Short-term plasticity for COM afferents was similar across target neurons at 10 Hz, but differed somewhat in the second half of trains delivered at 20 Hz (p ranging from 0.302 to 0.064 and *d* ranging from 0.32 to 0.61 over the final five EPSPs) or 40 Hz (p ranging from 0.124 to 0.037 and *d* ranging from 0.42 to 0.68 over the final four EPSPs). These data confirm that short-term plasticity is frequency dependent in COM and MDN afferents, but that synaptic dynamics of initial EPSPs are largely target-independent and similar across COM and MDN afferents.

To confirm that flash-evoked COM (**Figure 3C**) and MDN (**Figure 3D**) EPSPs are stable over at least 30 minutes of recording, we measured optically evoked EPSPs over time (n = 6 IT and 6 ET neurons for each afferent, not necessarily in pairs). After 10 minutes, a time point similar to when measurements of neuromodulation were made in experiments described below, average normalized first and second EPSP amplitudes were 1.00 ± 0.19 (p = 0.975, *d* = 0.05) and 1.02 ± 0.20 (p = 0.396, *d* = 0.08), respectively, and normalized CV and PPR were 1.13 ± 0.37 (p = 0.056, *d* = 0.20) and 1.03 ± 0.17 (p = 0.341, *d* = 0.08), respectively, relative to baseline values (n = 24; both afferents and target neurons combined). Our finding that polysynaptic MDN-driven EPSPs in ET neurons remain consistent over 30 minutes (**Figure 3D**) suggests generalized stability of cortical network dynamics over time in acute slices of the PL cortex.

### Neuromodulation of COM and MDN afferents

To test whether ACh and 5-HT differentially modulate COM and MDN synaptic transmission according to target neuron subtype, dual recordings were made from pairs of IT and ET target neurons while pairs of EPSPs were optically evoked at 20 Hz (15 s inter-trial intervals) in baseline conditions and after exposure to either 20 µM ACh (with 10 µM eserine; **Figure 4A-C**) or 40 µM 5-HT (**Figure 4D-F**). When COM EPSPs were challenged with ACh (n = 8 IT-ET pairs), optically evoked EPSPs were reversibly reduced to a similar extent in IT and ET target neurons (by ~85 ± 5%; **Table 5**), while CVs of EPSP amplitudes and PPRs were reversibly increased in both target neuron subtypes (**Figure 4A, B**; **Table 5**). These data suggest that Ach acutely suppresses glutamate release at COM synapses in a target-independent manner.

**Table 5.**
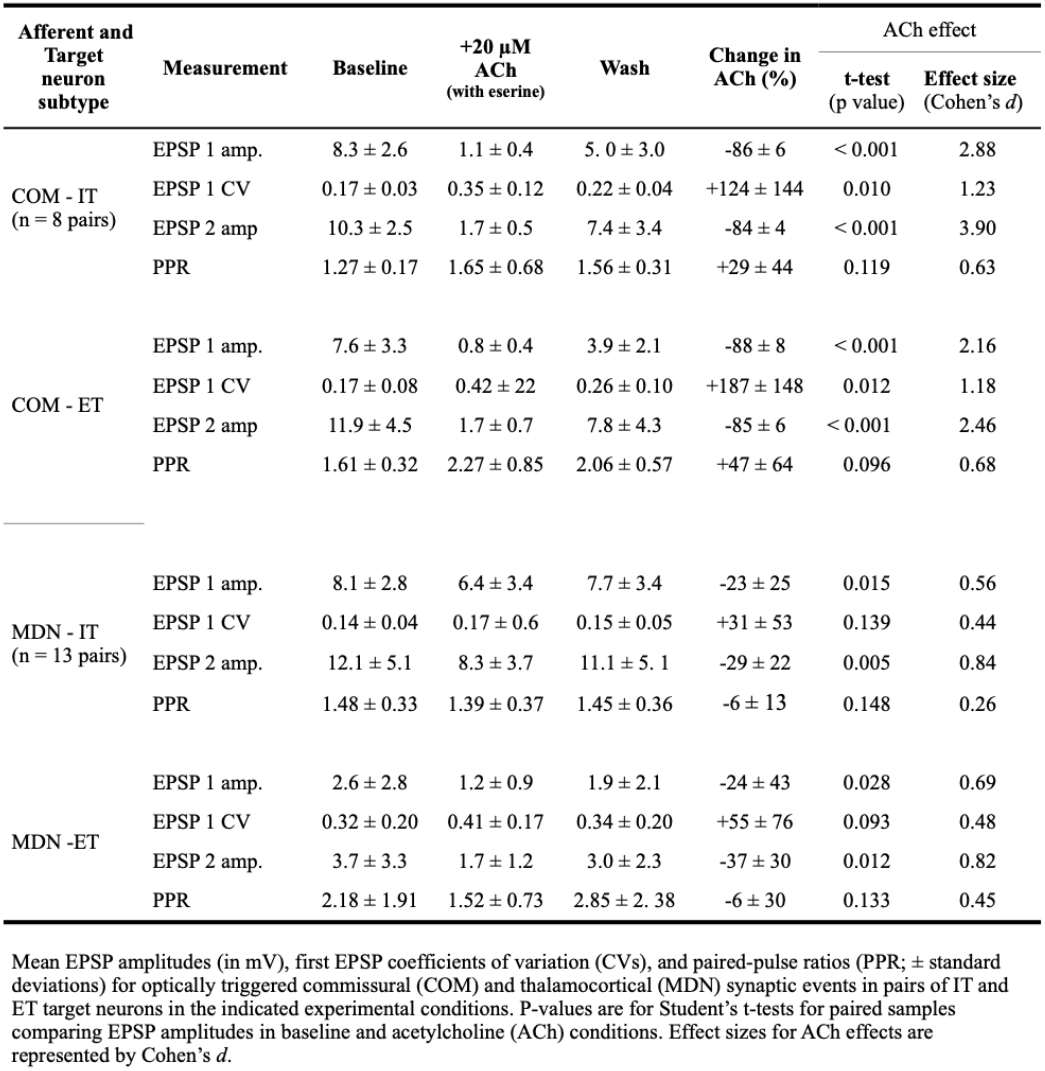
Cholinergic neuromodulation of COM and MDN afferents to IT and ET neurons.

| Afferent and Target neuron subtype | Measurement | Baseline | +20 μM ACh (with eserine) | Wash | Change in ACh (%) | ACh effect | |
| --- | --- | --- | --- | --- | --- | --- | --- |
|  |  |  |  |  |  | t-test (p value) | Effect size (Cohen's d) |
| COM - IT (n = 8 pairs) | EPSP 1 amp. | $8.3 \pm 2.6$ | $1.1 \pm 0.4$ | $5.0 \pm 3.0$ | $-86 \pm 6$ | < 0.001 | 2.88 |
| | EPSP 1 CV | $0.17 \pm 0.03$ | $0.35 \pm 0.12$ | $0.22 \pm 0.04$ | $+124 \pm 144$ | 0.010 | 1.23 |
| | EPSP 2 amp | $10.3 \pm 2.5$ | $1.7 \pm 0.5$ | $7.4 \pm 3.4$ | $-84 \pm 4$ | < 0.001 | 3.90 |
| | PPR | $1.27 \pm 0.17$ | $1.65 \pm 0.68$ | $1.56 \pm 0.31$ | $+29 \pm 44$ | 0.119 | 0.63 |
| COM - ET | EPSP 1 amp. | $7.6 \pm 3.3$ | $0.8 \pm 0.4$ | $3.9 \pm 2.1$ | $-88 \pm 8$ | < 0.001 | 2.16 |
| | EPSP 1 CV | $0.17 \pm 0.08$ | $0.42 \pm 0.22$ | $0.26 \pm 0.10$ | $+187 \pm 148$ | 0.012 | 1.18 |
| | EPSP 2 amp | $11.9 \pm 4.5$ | $1.7 \pm 0.7$ | $7.8 \pm 4.3$ | $-85 \pm 6$ | < 0.001 | 2.46 |
| | PPR | $1.61 \pm 0.32$ | $2.27 \pm 0.85$ | $2.06 \pm 0.57$ | $+47 \pm 64$ | 0.096 | 0.68 |
| MDN - IT (n = 13 pairs) | EPSP 1 amp. | $8.1 \pm 2.8$ | $6.4 \pm 3.4$ | $7.7 \pm 3.4$ | $-23 \pm 25$ | 0.015 | 0.56 |
| | EPSP 1 CV | $0.14 \pm 0.04$ | $0.17 \pm 0.6$ | $0.15 \pm 0.05$ | $+31 \pm 53$ | 0.139 | 0.44 |
| | EPSP 2 amp. | $12.1 \pm 5.1$ | $8.3 \pm 3.7$ | $11.1 \pm 5.1$ | $-29 \pm 22$ | 0.005 | 0.84 |
| | PPR | $1.48 \pm 0.33$ | $1.39 \pm 0.37$ | $1.45 \pm 0.36$ | $-6 \pm 13$ | 0.148 | 0.26 |
| MDN - ET | EPSP 1 amp. | $2.6 \pm 2.8$ | $1.2 \pm 0.9$ | $1.9 \pm 2.1$ | $-24 \pm 43$ | 0.028 | 0.69 |
| | EPSP 1 CV | $0.32 \pm 0.20$ | $0.41 \pm 0.17$ | $0.34 \pm 0.20$ | $+55 \pm 76$ | 0.093 | 0.48 |
| | EPSP 2 amp. | $3.7 \pm 3.3$ | $1.7 \pm 1.2$ | $3.0 \pm 2.3$ | $-37 \pm 30$ | 0.012 | 0.82 |
| | PPR | $2.18 \pm 1.91$ | $1.52 \pm 0.73$ | $2.85 \pm 2.38$ | $-6 \pm 30$ | 0.133 | 0.45 |

Mean EPSP amplitudes (in mV), first EPSP coefficients of variation (CVs), and paired-pulse ratios (PPR; ± standard deviations) for optically triggered commissural (COM) and thalamocortical (MDN) synaptic events in pairs of IT and ET target neurons in the indicated experimental conditions. P-values are for Student's t-tests for paired samples comparing EPSP amplitudes in baseline and acetylcholine (ACh) conditions. Effect sizes for ACh effects are represented by Cohen's $d$ .

**Figure 4.**
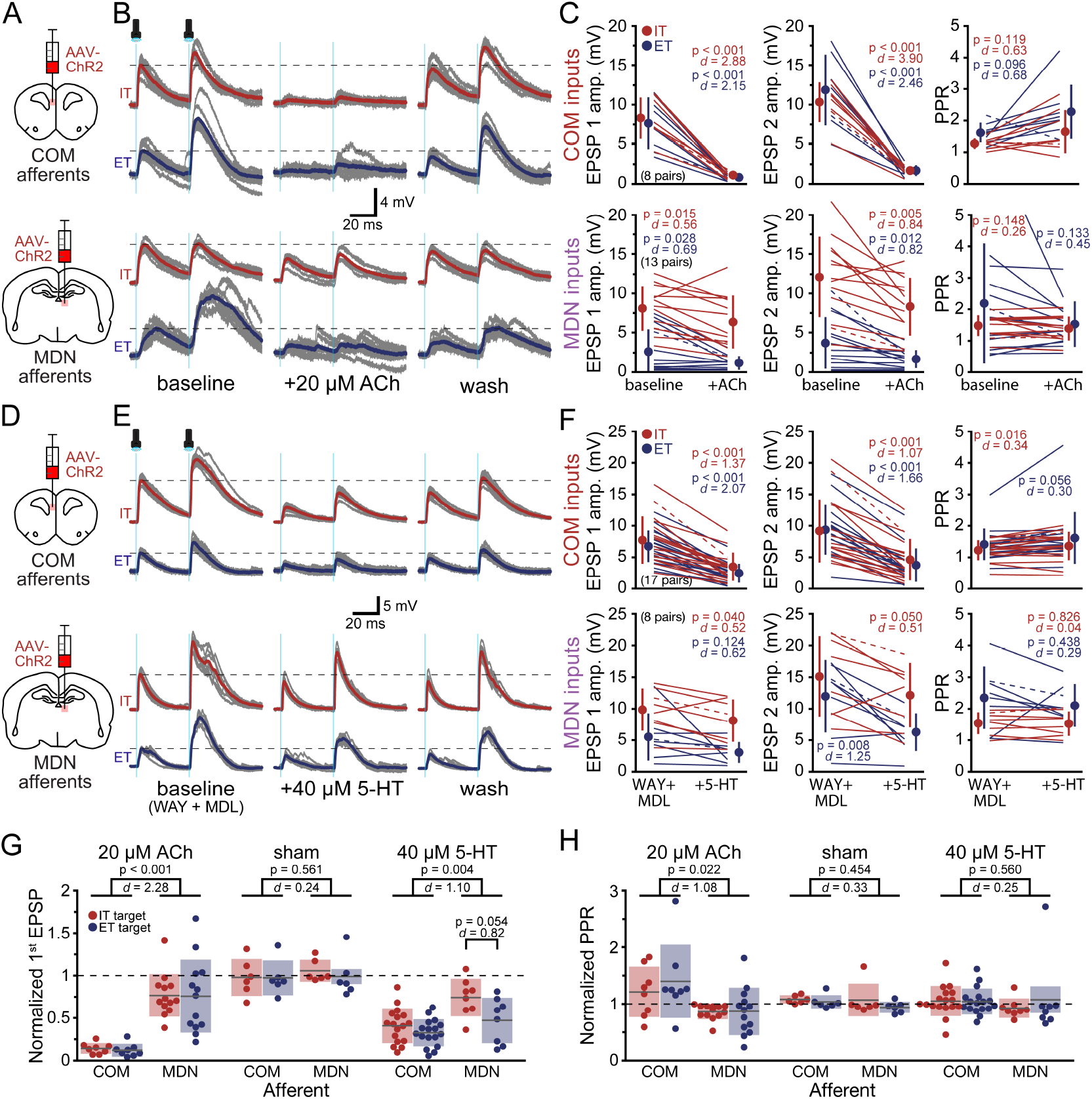
Cholinergic and serotonergic modulation of excitatory afferents in the prelimbic cortex. ***A***, Diagrams of approach for delivering ChR2-mCherry to commissural (COM; top) and thalamocortical (MDN; bottom) afferents to the prelimbic cortex. ***B***, Ten consecutive responses (grey) and their averaged responses (colored thick traces) in pairs of IT (red) and ET (blue) target neurons following activation of COM (top) or MDN (bottom) afferents in baseline conditions (left), after application of 20 µM ACh (middle), and after 15 - 20 minutes of wash (right). ***C***, Comparisons of EPSP amplitudes (left, middle) and paired-pulse ratios (PPR; right) in IT (red) and ET (blue) target neurons in baseline conditions and in the presence of ACh. Dashed lines indicate data from traces shown in B. Mean values and standard deviations are offset with filled symbols. P-values for Student’s t-tests and effect sizes (Cohen’s *d*) are color-coded to neuron subtype. ***D***-***F***, Similar to ***A-C***, but for application of 40 µM 5-HT. For 5-HT experiments, baseline conditions included WAY 100635 (30 nM) and MDL 11939 (500 nM) to block postsynaptic 5-HT receptors. Note that for MDN inputs, responses in ET neurons are largely polysynaptic (see main text). ***G***-***H***, Comparisons of treatment effects on first EPSP amplitudes and paired-pulse ratios for COM (***G***) and MDN (***H***) inputs to IT and ET neurons. Grey lines indicate means, and shaded areas denote standard deviations. The only large target-dependent effect was for serotonergic effects on MDN afferents (shown).

ACh had more modest effects on optically evoked MDN EPSPs (n = 13 IT-ET pairs; **Figure 4B, C**), with the first of two EPSPs being reduced by only ~25% (**Table 5**) and the second EPSPs being reduced by slightly larger amounts, leaving CVs and PPRs relatively unchanged in the presence of ACh (**Figure 4B, C**; **Table 5**). When cholinergic effects were compared across afferents, COM EPSPs in IT and ET target neurons were more sensitive to ACh than MDN inputs to IT neurons (amplitude: p < 0.001, *d* = 2.28; CV: p = 0.001, *d* = 1.27; **Figure 4G**; **Table 5**), and only COM inputs demonstrated increased PPRs in the presence of ACh (**Figure 4H**; **Table 5**). Responses to ACh in both afferents were larger than changes observed over time in sham experiments in which COM and MDN EPSPs were measured at similar time points but without ACh application (n = 18; see **Figure 3C, D**, and **Figure 4G, H**). When comparing ACh effects to sham experiments in COM afferents, p < 0.001 for both amplitude and CV, and *d* = 5.13 and 1.56, respectively for amplitude and CV. Comparing MDN inputs treated with ACh (IT targets only; n = 13) to MDN sham experiments (IT targets only; n = 6), p = 0.301 and *d* = 0.43. These data suggest that ACh preferentially suppresses glutamate release at COM terminals but does not have target-neuron-specific effects on synaptic transmission.

We next tested the impact of 5-HT on synaptic transmission at COM and MDN terminals. For these experiments, we first blocked postsynaptic 1A (with 30 nM WAY100635) and 2A (with 500 nM MDL11939) receptors to eliminate target-specific postsynaptic effects of 5-HT (e.g., Avesar and Gulledge, 2012) that might otherwise influence somatically recorded EPSPs. In these baseline conditions, optical activation of COM afferents generated robust EPSPs that were of similar amplitudes in IT and ET target neurons (n = 17 IT-ET pairs; **Figure 4E, F**; **Table 6**). Bath application of 40 µM 5-HT moderately reduced optically evoked COM EPSP amplitudes (by ~60%) and increased CVs (by ~60 to 80%) in both target neuron subtypes (**Table 6**), but had only modest and highly variable impact on PPRs (**Figure 4F**; **Table 6**). Serotonergic suppression of COM EPSPs was marginally more robust in ET target neurons than in IT targets (when comparing across IT and ET target neurons, p = 0.057 and 0.061, whereas *d* = 0.50 and 0.049, for 5-HT effects on the first and second EPSPs, respectively), and was long-lasting; 5-HT effects did not reverse within a 15-minute wash (**Table 6**).

**Table 6.** Serotonergic neuromodulation of COM and MDN afferents to IT and ET neurons.

| Afferent and Target neuron subtype (n) | Measurement | Baseline (MDL+WAY) | +40 μM 5-HT | Wash | Change in 5-HT (%) | 5-HT effect | |
| --- | --- | --- | --- | --- | --- | --- | --- |
|  |  |  |  |  |  | t-test (p value) | Effect size (Cohen's d) |
| COM - IT (n = 17 pairs) | EPSP 1 amp. | 7.7 ± 3.8 | 3.5 ± 2.2 | 4.6 ± 3.7 | -55 ± 18 | < 0.001 | 1.37 |
| | EPSP 1 CV | 0.17 ± 0.06 | 0.28 ± 0.11 | 0.25 ± 0.11 | +82 ± 72 | < 0.001 | 1.11 |
| | EPSP 2 amp. | 9.2 ± 5.0 | 4.6 ± 3.3 | 5.2 ± 4.3 | -52 ± 18 | < 0.001 | 1.07 |
| | PPR | 1.22 ± 0.32 | 1.35 ± 0.45 | 1.18 ± 0.43 | +9 ± 18 | 0.016 | 0.34 |
| COM - ET | EPSP 1 amp. | 6.7 ± 2.5 | 2.5 ± 1.5 | 3.0 ± 1.6 | -65 ± 13 | < 0.001 | 2.07 |
| | EPSP 1 CV | 0.17 ± 0.05 | 0.26 ± 0.10 | 0.23 ± 0.07 | +61 ± 78 | 0.002 | 0.91 |
| | EPSP 2 amp. | 9.3 ± 4.0 | 3.7 ± 2.7 | 4.7 ± 2.8 | -61 ± 14 | < 0.001 | 1.66 |
| | PPR | 1.41 ± 0.50 | 1.61 ± 0.82 | 1.70 ± 0.98 | +12 ± 18 | 0.056 | 0.30 |
| MDN - IT (n = 8 pairs) | EPSP 1 amp. | 9.8 ± 3.3 | 8.1 ± 3.4 | 7.7 ± 3.4 | -26 ± 22 | 0.040 | 0.52 |
| | EPSP 1 CV | 0.10 ± 0.03 | 0.12 ± 0.05 | 0.13 ± 0.05 | +25 ± 63 | 0.389 | 0.32 |
| | EPSP 2 amp. | 15.1 ± 6.3 | 12.1 ± 5.1 | 11.1 ± 5.1 | -27 ± 19 | 0.050 | 0.51 |
| | PPR | 1.54 ± 0.35 | 0.83 ± 0.04 | 1.45 ± 0.36 | +0.3 ± 17 | 0.826 | 0.04 |
| MDN - ET | EPSP 1 amp. | 5.5 ± 3.7 | 3.0 ± 1.7 | 5.1 ± 1.9 | -53 ± 27 | 0.124 | 0.62 |
| | EPSP 1 CV | 0.18 ± 0.09 | 0.21 ± 0.08 | 0.18 ± 0.08 | +27 ± 61 | 0.281 | 0.35 |
| | EPSP 2 amp. | 12.0 ± 5.7 | 6.3 ± 3.0 | 10.6 ± 4.5 | -57 ± 19 | 0.008 | 1.25 |
| | PPR | 2.34 ± 0.99 | 2.10 ± 0.68 | 2.08 ± 0.86 | +8 ± 67 | 0.438 | 0.29 |

Mean EPSP amplitudes (in mV), first EPSP coefficients of variation (CVs), and paired-pulse ratios (PPR; ± standard deviations) for optically triggered commissural (COM) and thalamocortical (MDN) synaptic events in pairs of IT and ET target neurons in the indicated experimental conditions. P-values are for Student's t-tests for paired samples comparing EPSP amplitudes in baseline and serotonin (5-HT) conditions. Effect sizes for ACh effects are represented by Cohen's *d*.

Serotonergic effects on MDN EPSPs were also modest (**Figure 4E, F**; **Table 6**), particularly in IT target neurons, where MDN EPSPs were suppressed by only ~25% relative to the ~55% reduction of EPSP amplitudes in ET neurons (n = 8 IT-ET pairs; p = 0.054 and 0.004, *d* = 0.82 and 1.48, for first and second EPSPs, respectively, when comparing across target neuron subtypes; **Figure 4G**). For both afferents, serotonergic effects were larger than those observed in sham experiments in which there was almost no change in the amplitude of EPSPs measured at a similar time point (see above and **Figure 4G**). Finally, 5-HT had relatively little impact on CVs and PPRs in both ET and IT target neurons (**Figure 4F, H**; **Table 6**). These data suggest that 5-HT, like ACh, may preferentially suppress glutamate release at COM terminals while leaving MDN synaptic transmission largely unaffected.

### Neuromodulation of monosynaptic inputs to IT and ET neurons

The data described above suggest that both ACh and 5-HT may preferentially suppress glutamatergic synaptic transmission at COM, relative to MDN, terminals in the PL cortex. However, activation of excitatory afferents may trigger network activity (i.e., polysynaptic transmission) in the local cortical circuit that might contribute to the effects of ACh or 5-HT on synaptic transmission. For instance, since MDN-evoked responses in ET neurons are largely polysynaptic (**Figure 2**), neuromodulation of MDN EPSPs in ET neurons may reflect changes elsewhere in the local network rather than direct effects at MDN terminals.

To test whether ACh and/or 5-HT suppresses monosynaptic transmission from COM and MDN afferents, we conducted additional experiments in the presence of 1 µM TTX and 100 µM 4-AP. ACh suppressed isolated monosynaptic COM EPSPs by ~80 ± 12%, with negligible changes in PPR (n = 6 IT and 4 ET neurons, not necessarily in pairs; **Figure 5A, B**; **Table 7**). Alternatively, when monosynaptic MDN inputs were activated, EPSPs were largely absent in ET neurons (n = 2) and those occurring in IT target neurons were relatively insensitive to ACh (n = 7; **Figure 5C, D**; **Table 7**). When comparing cholinergic modulation of monosynaptic COM-(all target neurons) and MDN-(IT targets only) evoked EPSPs, the effect of ACh on initial EPSP amplitudes was ~7-fold greater for COM afferents (at −78 ± 10%) than for MDN afferents (−11 ± 15%; n = 17, p < 0.001, *d* = 2.02, when comparing across afferents). These data suggest that ACh selectively suppresses glutamate release from COM synaptic terminals in the PL cortex. Afferent-specific neuromodulation of monosynaptic transmission also points to a presynaptic locus of cholinergic effects, as postsynaptic effects of ACh on synaptic transmission are expected to be largely similar across afferents.

**Table 7.** Neuromodulation of monosynaptic transmission in IT and ET target neurons.

| Afferent | Measure | Baseline<br>(TTX+4-AP) | +ACh | Wash | % Change<br>in ACh | t-test<br>(p value) | Effect size<br>(Cohen's *d*) |
| --- | --- | --- | --- | --- | --- | --- | --- |
| COM<br>(n = 10) | EPSP 1 amp. | 10.5 ± 4.5 | 2.6 ± 2.0 | 8.0 ± 4.5 | -78 ± 10 | < 0.001 | 2.29 |
|  | EPSP 1 CV | 0.20 ± 0.07 | 0.38 ± 0.20 | 0.20 ± 0.09 | +92 ± 87 | 0.015 | 0.94 |
|  | EPSP 2 amp. | 14.8 ± 12.1 | 4.1 ± 4.2 | 10.3 ± 9.1 | -77 ± 13 | 0.002 | 1.19 |
|  | PPR | 1.35 ± 0.64 | 1.45 ± 0.75 | 1.23 ± 0.71 | +7 ± 36 | 0.402 | 0.15 |
| MDN<br>(n = 7,<br>IT only) | EPSP 1 amp. | 9.8 ± 3.8 | 8.6 ± 2.9 | 11.4 ± 3.9 | -11 ± 15 | 0.043 | 0.38 |
|  | EPSP 1 CV | 0.13 ± 0.05 | 0.13 ± 0.04 | 0.15 ± 0.11 | +17 ± 64 | 0.903 | 0.05 |
|  | EPSP 2 amp. | 8.5 ± 4.2 | 7.7 ± 4.3 | 10.2 ± 5.8 | -7 ± 23 | 0.359 | 0.18 |
|  | PPR | 0.87 ± 0.25 | 0.91 ± 0.30 | 0.86 ± 0.25 | +4 ± 20 | 0.558 | 0.15 |
| Afferent | Measure | Baseline<br>(TTX+4-AP) | +5-HT | Wash | % Change<br>in 5-HT | t-test<br>(P value) | Effect size<br>(Cohen's *d*) |
| COM<br>(n = 14) | EPSP 1 amp. | 10.5 ± 3.4 | 6.1 ± 2.8 | * | -44 ± 16 | < 0.001 | 1.41 |
|  | EPSP 1 CV | 0.19 ± 0.09 | 0.20 ± 0.07 | * | +33 ± 87 | 0.687 | 0.13 |
|  | EPSP 2 amp. | 11.7 ± 4.5 | 7.1 ± 3.6 | * | -41 ± 19 | 0.002 | 1.13 |
|  | PPR | 1.13 ± 0.32 | 1.19 ± 0.44 | * | +4 ± 25 | 0.464 | 0.15 |
| MDN<br>(n = 11,<br>IT only) | EPSP 1 amp. | 11.6 ± 4.1 | 13.7 ± 9.9 | * | +12 ± 58 | 0.391 | 0.29 |
|  | EPSP 1 CV | 0.19 ± 0.21 | 0.16 ± 0.11 | * | +54 ± 216 | 0.866 | 0.12 |
|  | EPSP 2 amp. | 14.1 ± 8.8 | 13.3 ± 8.9 | * | -7 ± 35 | 0.622 | 0.09 |
|  | PPR | 1.18 ± 0.55 | 0.99 ± 0.37 | * | -9 ± 26 | 0.296 | 0.40 |

Mean EPSP amplitudes (in mV), first EPSP coefficients of variation (CVs), and paired-pulse ratios (PPR; ± standard deviations) for isolated monosynaptic commissural (COM) and thalamocortical (MDN) synaptic events under baseline and neuromodulatory conditions. For COM afferents, IT and ET targets were combined. For MDN inputs, data are presented from IT targets only. Baseline EPSPs were measured in the presence of 1 μM tetrodotoxin (TTX) and 100 μM 4-aminopyridine (4-AP). \*For 5-HT experiments, not all cells were allowed a full wash in TTX+4-AP. P-values are for Student's t-tests for paired samples, while effect sizes are represented by Cohen's *d*.

**Figure 5.**
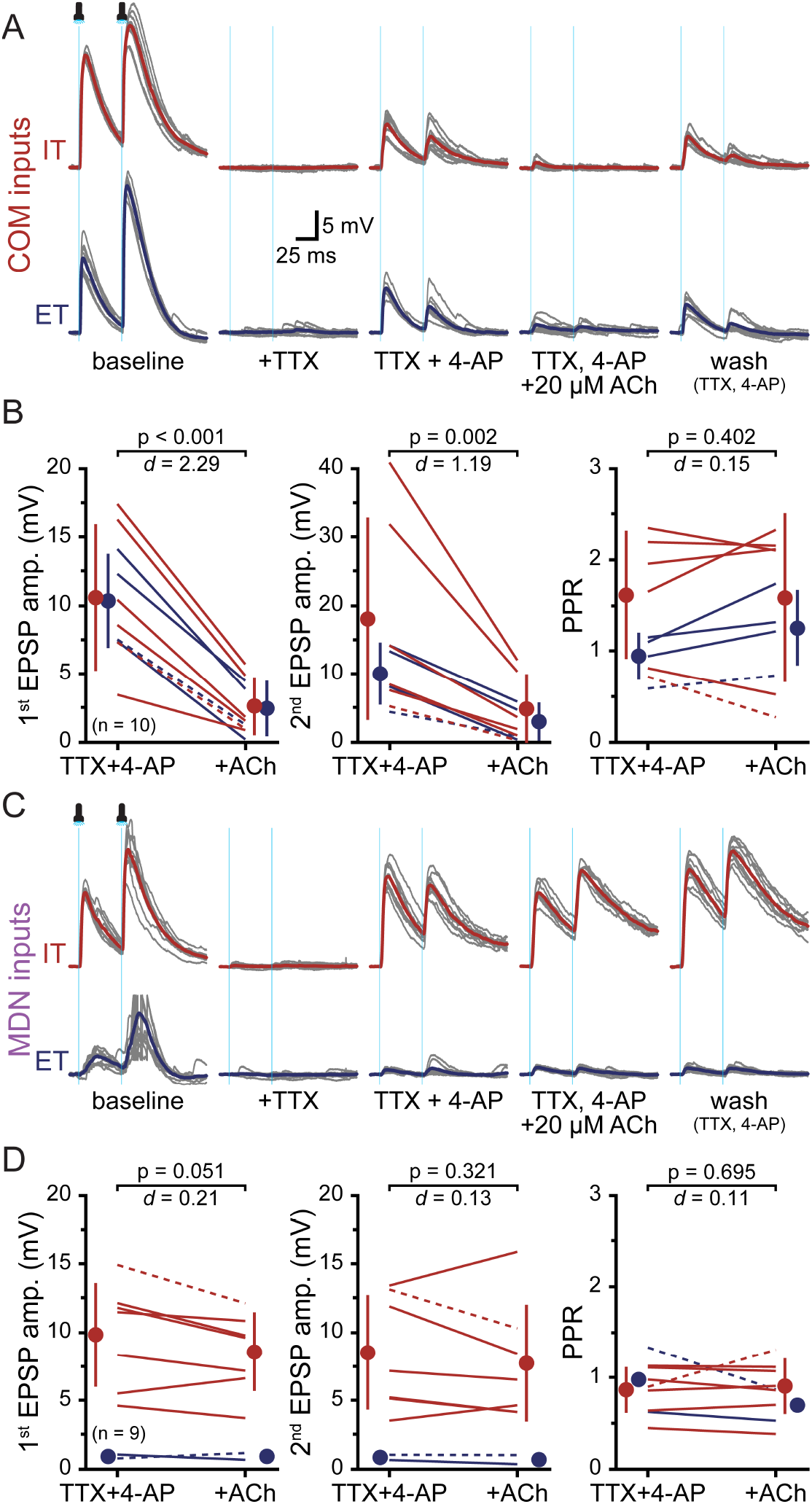
Selective cholinergic suppression of glutamate release from monosynaptic commissural (COM) afferents to IT and ET neurons in prelimbic cortex. ***A***, Ten consecutive optically activated COM EPSP pairs (grey traces) and their averaged responses (colored traces) in a pair of IT (red) and ET (blue) neurons in baseline conditions (left), after sequential and additive applications of 1 µM TTX, 100 µM 4-AP, and 20 µM Ach (with 10 µM eserine; middle traces), and after wash in TTX + 4-AP (right). ***B***, Plots of the effects of 20 µM ACh on COM EPSP amplitudes (left and middle) and paired-pulse ratios (PPR; right). Colored lines denote data from individual IT (red) or ET (blue) neurons, with dashed lines indicating data for the neurons shown in *A*. Mean values and standard deviations are offset with filled symbols. ***C***, Ten consecutive responses (grey) and their averages (colored traces) for optical activation of thalamic (MDN) inputs to a pair of IT (red) and ET (blue) neurons in the indicated experimental conditions. Note the lack of monosynaptic MDN input to the ET neuron. ***D***, Plots of the effects of 20 µM ACh on MDN EPSP amplitudes (left and middle) and paired-pulse ratios (PPR; right). Colored lines denote data from individual IT (red) or ET (blue) neurons, with dashed lines indicating data for the neurons shown in ***C***. Mean values and standard deviations are offset with filled symbols. P-values (Student’s t-tests for paired samples) and effect sizes (Cohen’s *d*) in ***B*** and ***D*** are calculated from all neurons (for IT-only results for MDN inputs, see **Table 7**).

5-HT also suppressed monosynaptic COM EPSPs, albeit to a lesser extent than observed with ACh. In the absence of blockers of 1A and 2A receptors, the addition of 40 µM 5-HT reduced isolated monosynaptic COM EPSPs by ~40 - 45% (n = 7 IT and 7 ET neurons, not necessarily pairs; **Figure 6A, B**; **Table 7**), but failed to suppress monosynaptic MDN EPSPs in IT target neurons (n = 11; **Figure 6C, D**; **Table 7**). When comparing the effects of 5-HT on first EPSP amplitude across afferents, 5-HT had a much larger effect on COM afferents relative to MDN inputs (n = 25; p = 0.010, *d* = 1.39).

**Figure 6.**
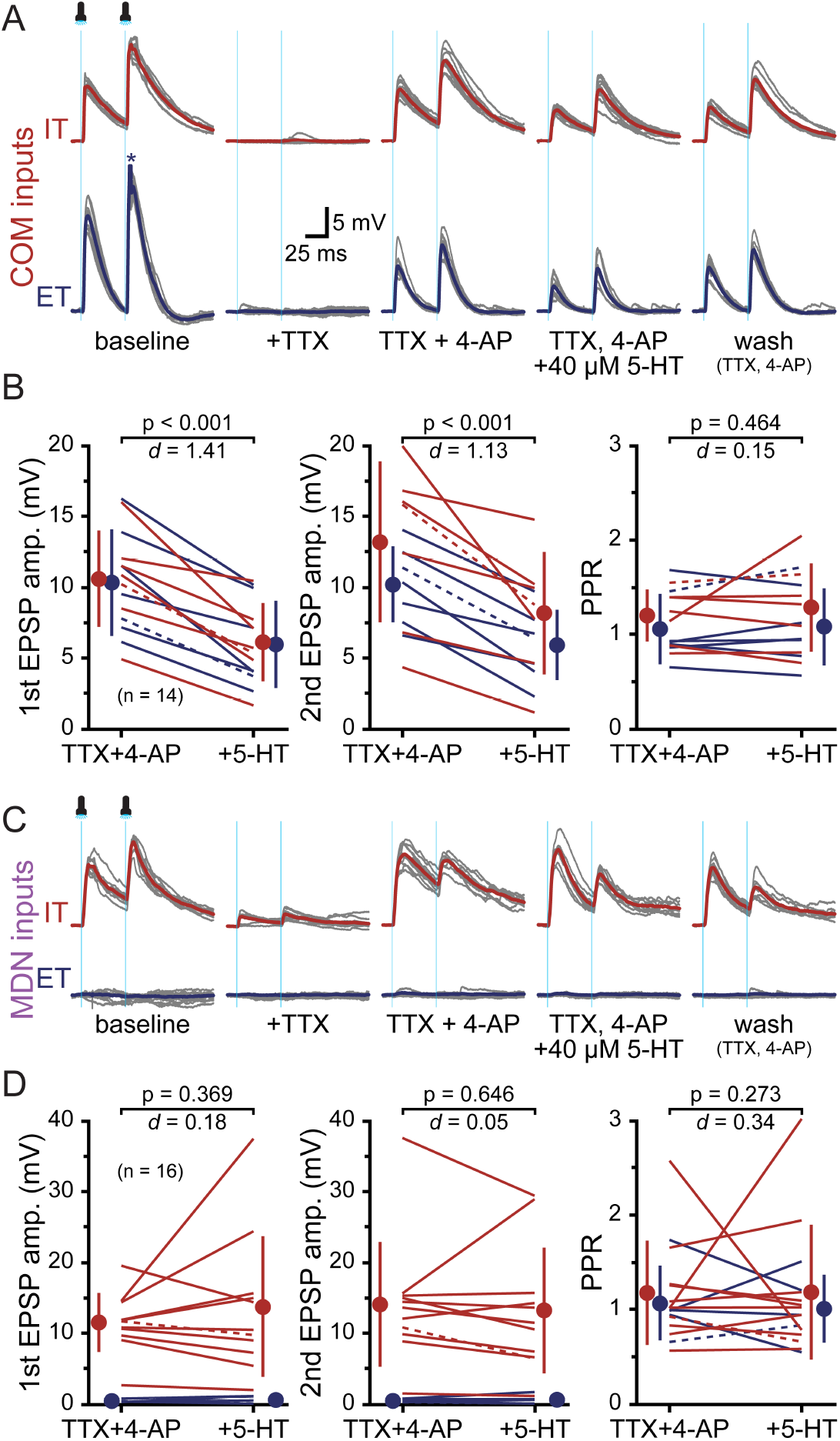
Selective serotonergic suppression of glutamate release from monosynaptic commissural (COM) afferents to IT and ET neurons in prelimbic cortex. ***A***, Ten consecutive optically activated COM EPSP pairs (grey traces) and their averaged responses (colored traces) in a pair of IT (red) and ET (blue) neurons in baseline conditions (left), after sequential and additive applications of 1 µM TTX, 100 µM 4-AP, and 40 µM 5-HT (middle traces), and after wash in TTX + 4-AP (right). ***B***, Plots of the effects of 40 µM 5-HT on COM EPSP amplitudes (left and middle) and paired-pulse ratios (PPR; right). Colored lines denote data from individual IT (red) or ET (blue) neurons, with dashed lines indicating data for the neurons shown in *A*.Mean values and standard deviations are offset with filled symbols. ***C***, Ten consecutive EPSP traces (grey) and their averages (colored traces) for thalamic (MDN) inputs to a pair of IT (red) and ET (blue) neurons in the indicated experimental conditions. ***D***, Plots of the effects of 40 µM 5-HT on MDN EPSP amplitudes (left and middle) and paired-pulse ratios (PPR; right). Colored lines denote data from individual IT (red) or ET (blue) neurons, with dashed lines indicating data for the neurons shown in ***C***. Mean values and standard deviations are offset with filled symbols. P-values (Student’s t-tests for paired samples) and effect sizes (Cohen’s *d*) in ***B*** and ***D*** are calculated from all neurons (for IT-only results for MDN inputs, see **Table 7**).

In the presence of TTX and 4-AP, application of ACh or 5-HT produced little change in PPRs or CVs for either COM or MDN inputs (**Figure 5D, 6D**; **Table 7**). This likely reflects differences in presynaptic release probability that occur when synaptic transmission is triggered directly by ChR2, rather than with action potentials (see, for instance, Cruikshank et al., 2010), and likely also explains the lower overall baseline PPRs measured in the presence of TTX and 4-AP (see also **Figure 2C**). To our knowledge, this is the first report of cholinergic or serotonergic modulation of isolated monosynaptic optically triggered synaptic transmission in the cortex, with our results suggesting that both ACh and 5-HT regulate glutamate release selectively at COM terminals while having little impact on MDN synaptic transmission.

### Concentration-dependent neuromodulation of COM input to IT and ET neurons

We next measured the effects of ACh and 5-HT on COM synaptic transmission across a range of transmitter concentrations. For these experiments, recordings were made of COM-evoked EPSPs (2 flashes at 20 Hz at 15 s intervals) in IT and ET target neurons (not necessarily in pairs) in tissue exposed sequentially to increasing concentrations of modulatory transmitters. For cholinergic modulation, COM EPSPs were measured in two groups of neurons in baseline conditions, across three increasing concentrations of ACh applied for 7 minutes each, and after an additional 15 minutes of wash.

The first group (n = 7 IT and 10 ET neurons) was exposed sequentially to eserine alone (10 µM), eserine with 500 nM ACh, and eserine with 20 µM ACh, followed by a 15-minute wash period (**Figure 7A**). The second group (n = 8 IT and 7 ET neurons) experienced sequential applications of 1, 10, and 40 µM ACh, all in the presence of 10 µM eserine, before a 15-minute wash period (**Figure 7A**). Bath application of eserine alone was sufficient to reduce EPSP amplitudes (by an average of 39 ± 14% and 32 ± 3.3% for the first and second EPSPs, respectively; n = 17; p < 0.001, *d* ≥ 1.15 for each) and increase PPRs (by 13 ± 14%; p = 0.002, *d* = 0.81), suggesting an endogenous level of ACh in the absence of acetylcholinesterase that is sufficient to modulate synaptic transmission at COM afferents (**Figure 7B**). We found that increasing ACh concentrations progressively reduced EPSP amplitudes and increased PPRs, saturating at about 10 µM ACh in the presence of eserine with an 88 ± 7% reduction in the first EPSP (**Figure 7B**). Further, sequential cholinergic manipulations steadily increased the CV of initial EPSP amplitudes. In group one, CVs increased from a baseline average of 0.18 ± 0.10 (n = 15) to values of 0.23 ± 0.07 (in eserine only; p = 0.053, *d* = 0.55, relative to baseline), 0.31 ± 0.14 (500 nM ACh in eserine; p < 0.001, *d* = 1.08), and 0.31 ± 0.07 (20 µM ACh in eserine; p < 0.001, *d* = 1.26). For group 2 neurons (n = 15), CVs increased from baseline values of 0.17 ± 0.07 to 0.37 ± 0.15 (1 µM ACh in eserine), 0.40 ± 0.21 (10 µM ACh in eserine), and 0.51 ± 0.21 (40 µM ACh in eserine), with all p values below 0.001 and all effect sizes (Cohen’s *d*) at or above 1.30. Fitting a Hill equation to the mean amplitudes of initial EPSPs across experimental conditions (**Figure 7A**, inset; see Methods) allowed us to estimate the endogenous (i.e., eserine-revealed) ACh concentration to be ~139 nM, and using that value we calculated the IC50 for cholinergic suppression of COM EPSPs to be 177 ± 15 nM (mean ± SD).

**Figure 7.**
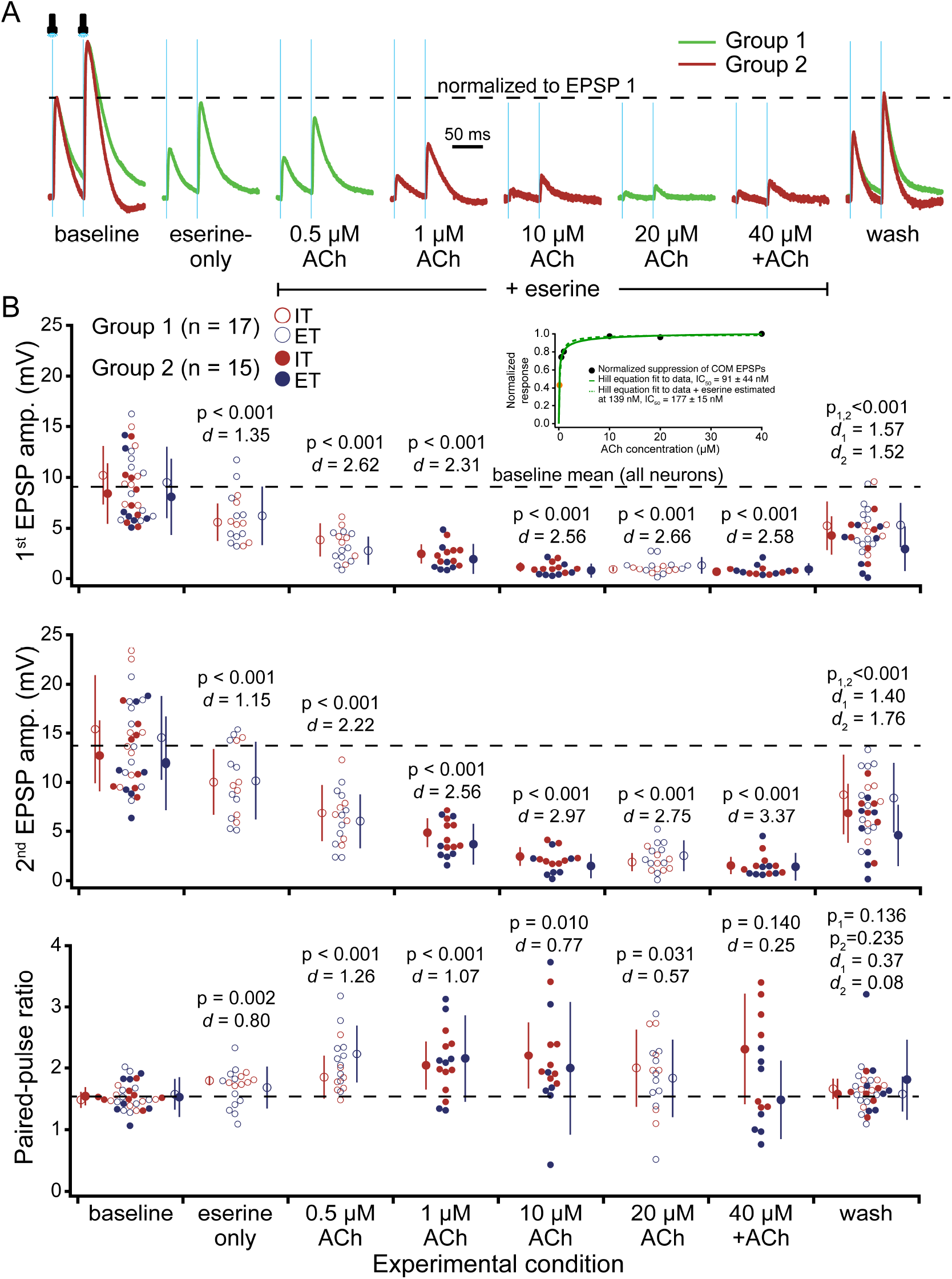
(facing page) Dose-dependence of cholinergic suppression of excitatory transmission at commissural (COM) afferents. ***A***, Averages of ten consecutive light-evoked COM EPSPs (20 Hz; normalized to baseline EPSP amplitudes) in an IT neuron from “Group 1” and an ET neuron from “Group 2”. After recording baseline COM EPSPs, IT and ET neurons in Group 1 received three consecutive treatments consisting of eserine alone (10 µM), eserine with 0.5 µM ACh, and eserine with 20 µM ACh, followed by 10 or more minutes of wash. Group 2 IT and ET neurons were exposed to eserine combined with three increasing doses of ACh (1, 10, and 40 µM). Each dose was applied for 7 minutes. ***B***, Plots of EPSP amplitudes (top two graphs) and paired-pulse ratios (bottom graph) for each neuron in the various experimental conditions (as measured from averages of final 10 responses per experimental condition) for IT (red) and ET (blue) neurons in group 1 (open symbols) and Group 2 (closed symbols). Mean response amplitudes and standard deviations for each condition indicated by larger symbols to the left (IT; red) and right (ET; blue) of each data set. Inset is a plot of normalized responses vs ACh concentrations fit with a Hill equation for raw data (solid green line) and after estimating the eserine-only concentration at 139 nM (orange data point and dashed green line). P-values and Cohen’s *d* for each experimental condition are relative to baseline measurements for the given group. Neurons were not necessarily recorded in pairs.

A similar set of experiments was conducted using increasing concentrations of 5-HT (in the absence of any 5-HT receptor antagonists; **Figure 8**). For these experiments, group 1 neurons (n = 11 IT and 7 ET) experienced sequential applications of 100 nM, 500 nM and 2 µM 5-HT while group 2 neurons (n = 12 IT and 5 ET) experienced 10, 40, and 100 µM 5-HT (**Figure 8A**). Weak suppression of COM EPSPs was observed at 100 nM, the lowest dose tested, where the first and second EPSPs were reduced by 20 ± 13% and 17 ± 10%, respectively (n = 18), but was robust at 2 µM 5-HT, where the first and second EPSPs were reduced by 66 ± 21% and 64 ± 15%, respectively (**Figure 8B**). Similarly, CVs of first EPSP amplitudes (baseline values of 0.16 ± 0.06 and 0.16 ± 0.05, for groups 1 and 2, respectively) were markedly larger at and beyond 2 µM 5-HT. For group 1 responses, CVs rose from a baseline of 0.16 ± 06 (n = 19) to 0.17 ± 0.05 (p = 0.370, *d* = 0.21), 0.21 ± 0.07 (p = 0.009, *d* = 0.30), and 0.31 ± 0.11 (p < 0.001, *d* = 1.25) for sequential exposures to 100 nM, 500 nM, and 2 µM 5-HT, respectively. In group 2 neurons, CVs increased from baseline values of 0.16 ± 0.05 to 0.38 ± 0.18, 0.30 ± 11, and 0.32 ± 0.11 for sequential exposures to 10 µM, 40 µM, and 100 µM 5-HT, respectively (p values all below 0.001 with effect sizes ≥ 1.27). Fitting a Hill equation to the mean amplitudes for initial EPSPs across all concentrations indicated an IC50 of 216 ± 38 nM (**Figure 8B**).

**Figure 8.**
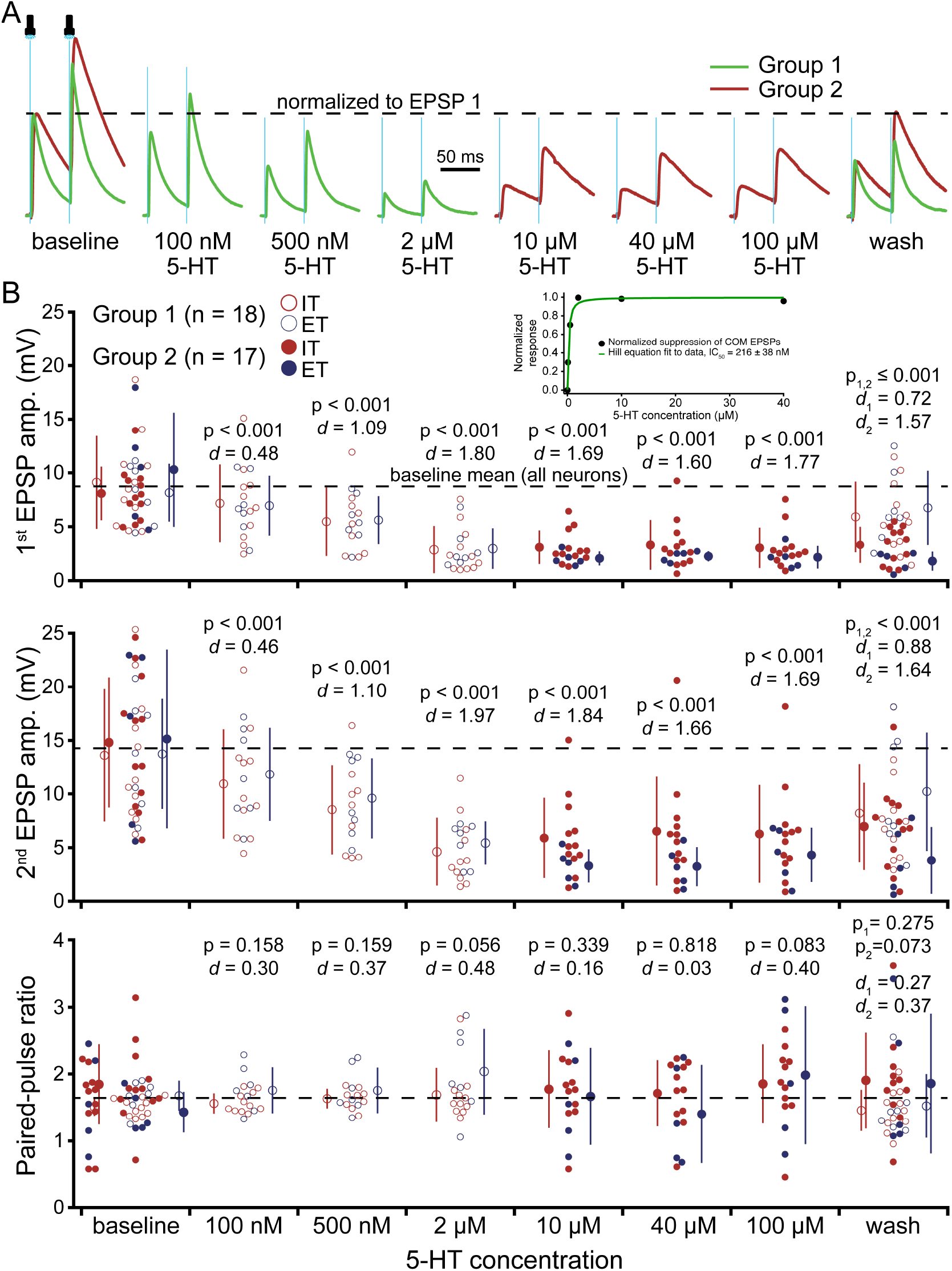
(facing page) Dose-dependence of serotonergic suppression of excitatory transmission at commissural (COM) afferents. ***A***, Averages of ten consecutive light-evoked COM EPSPs (20 Hz; normalized to baseline EPSP amplitudes) in two IT neurons (one from “Group 1” and one from “Group 2”). After recording baseline COM EPSPs, IT and ET neurons in Group 1 received three consecutively increasing doses of 5-HT (100 nM, 500 nM, and 2 µM) followed by 10 or more minutes of wash. Group 2 IT and ET neurons were sequentially exposed to 10, 40, and 100 µM 5-HT. ***B***, Plots of EPSP amplitudes (top two graphs) and paired-pulse ratios (bottom graph) for each neuron in the various experimental conditions (as measured from the averages of ten consecutive trials per condition) for IT (red) and ET (blue) neurons in group 1 (open symbols) and Group 2 (closed symbols). Mean response amplitudes and standard deviations for each experimental condition indicated by larger symbols to the left (IT; red) and right (ET; blue) of each data set. Inset is a plot of normalized responses vs 5-HT concentration fit with a Hill equation. Data from one Group 2 neuron with PPR values above 4 (4.8, 9.5, 4.6, and 6.4, respectively, for 10 µM, 40 µM, 100 µM, and wash) were excluded for calculations of mean and standard deviation and are not shown in plot. An additional pair of group 2 neurons was not exposed to 40 µM 5-HT. P-values and Cohen’s *d* for each experimental condition are relative to baseline measurements for the given group. Neurons were not necessarily recorded in pairs.

Interestingly, at 40 µM 5-HT, serotonergic suppression of the first (−64 ± 22%) and second (−64 ± 19%) COM EPSPs was only slightly greater in magnitude than the effects of 40 µM 5-HT in the presence of antagonists for 1A and 2A receptors (−60 ± 16%; p = 0.218, *d* = 0.43 for suppression of the first EPSP, and −57 ± 16%; p = 0.073, *d* = 0.64 for the second EPSPs; see **Figure 4** and **Table 6**, above), suggesting that 1A and 2A receptors are not major contributors to serotonergic modulation of glutamate release at COM afferents.

### Pharmacology of cholinergic modulation of COM afferents

We next determined the receptor subtypes mediating neuromodulation of synaptic transmission at COM terminals. Given that both ACh and 5-HT suppress, rather than enhance, COM transmission across a range of concentrations, we focused on the Gi/o-coupled metabotropic receptors that are most associated with inhibition of glutamate release in the cortex. In the case of ACh, metabotropic signaling is mediated by muscarinic ACh receptors (mAChRs), and so we first tested their broad involvement using atropine (1 µM), a non-specific mAChR antagonist. In the presence of atropine, pairs of COM EPSPs optically evoked at 20 Hz were 9.9 ± 2.4 mV and 15.1 ± 2.6 mV in amplitude for first and second EPSPs, respectively, and exhibited a PPR of 1.57 ± 0.26 (n = 8; **Figure 9A, B**; **Table 8**). The addition of 20 µM ACh for five minutes reduced the first and second EPSP amplitudes by ~10%, to 8.8 ± 2.2 mV and 13.5 ± 2.6 mV, respectively, but had minimal impact on PPR (1.55 ± 0.16; see **Figure 9B** for statistics). These modest effects were dramatically less than those observed in the absence of atropine (n = 8; see **Figure 4C**; **Table 5**) for both the first (n = 16; p < 0.001, *d* = 10.70, comparing experiments with and without atropine) and second (p < 0.001, *d* = 11.41) EPSPs, and for PPRs (p = 0.112, *d* = 0.90), suggesting that cholinergic modulation of COM synaptic transmission relies primarily on mAChRs.

**Table 8.** Pharmacology of cholinergic modulation of COM EPSPs.

| Drug condition | Measure | Baseline | +ACh | Wash | % Change in ACh | t-test (p value) | Effect size (Cohen's d) |
| --- | --- | --- | --- | --- | --- | --- | --- |
| 20 $\mu\text{M}$ ACh in 1 $\mu\text{M}$ atropine (muscarinic antagonist; $n = 8$) | EPSP 1 amp. | $9.9 \pm 2.4$ | $8.8 \pm 2.2$ | $8.3 \pm 2.2$ | $-10 \pm 8$ | 0.007 | 0.46 |
| | EPSP 1 CV | $0.13 \pm 0.04$ | $0.12 \pm 0.04$ | $0.15 \pm 0.05$ | $0 \pm 42$ | 0.684 | 0.15 |
| | EPSP 2 amp. | $15.1 \pm 2.6$ | $13.5 \pm 2.6$ | $13.2 \pm 3.1$ | $-11 \pm 8$ | 0.011 | 0.61 |
| | PPR | $1.57 \pm 0.26$ | $1.55 \pm 0.16$ | $1.98 \pm 0.47$ | $0 \pm 9$ | 0.778 | 0.07 |
| 1 $\mu\text{M}$ ACh alone ($n = 15$) | EPSP 1 amp. | $8.3 \pm 3.2$ | $3.6 \pm 0.9$ | $3.6 \pm 2.1$ | $-73 \pm 12$ | $< 0.001$ | 2.31 |
| | EPSP 1 CV | $0.18 \pm 0.07$ | $0.37 \pm 0.15$ | $0.24 \pm 0.09$ | $+137 \pm 128$ | 0.001 | 1.12 |
| | EPSP 2 amp. | $12.4 \pm 4.1$ | $4.3 \pm 1.8$ | $5.7 \pm 3.2$ | $-65 \pm 10$ | $< 0.001$ | 2.56 |
| | PPR | $1.54 \pm 0.24$ | $2.10 \pm 0.54$ | $1.71 \pm 0.50$ | $+38 \pm 36$ | 0.001 | 1.07 |
| 1 $\mu\text{M}$ ACh in 5 $\mu\text{M}$ AF-DX 116 (M2 antagonist; $n = 8$) | EPSP 1 amp. | $8.9 \pm 2.7$ | $8.9 \pm 2.7$ | $6.1 \pm 1.6$ | $-31 \pm 7$ | $< 0.001$ | 1.30 |
| | EPSP 1 CV | $0.14 \pm 0.03$ | $0.18 \pm 0.04$ | $0.23 \pm 0.08$ | $+37 \pm 42$ | 0.040 | 0.89 |
| | EPSP 2 amp. | $15.3 \pm 4.2$ | $11.9 \pm 3.6$ | $11.0 \pm 3.8$ | $-22 \pm 10$ | 0.001 | 0.87 |
| | PPR | $1.72 \pm 0.18$ | $1.95 \pm 0.29$ | $1.98 \pm 0.47$ | $+13 \pm 9$ | 0.006 | 0.94 |
| 1 $\mu\text{M}$ ACh in 5 $\mu\text{M}$ VU6028418 (M4 antagonist; $n = 15$) | EPSP 1 amp. | $7.2 \pm 2.5$ | $6.2 \pm 1.8$ | $6.6 \pm 2.5$ | $-12 \pm 12$ | 0.001 | 0.44 |
| | EPSP 1 CV | $0.18 \pm 0.06$ | $0.22 \pm 0.05$ | $0.22 \pm 0.08$ | $+33 \pm 49$ | 0.058 | 0.53 |
| | EPSP 2 amp. | $12.2 \pm 3.2$ | $10.5 \pm 2.7$ | $11.1 \pm 3.4$ | $-12 \pm 11$ | 0.001 | 0.54 |
| | PPR | $1.75 \pm 0.23$ | $1.72 \pm 0.21$ | $1.75 \pm 0.27$ | $+1 \pm 11$ | 0.826 | 0.05 |
| 1 $\mu\text{M}$ ACh in 3 $\mu\text{M}$ VU0467154 (M4 PAM; $n = 8$) | EPSP 1 amp. | $7.1 \pm 2.0$ | $1.0 \pm 0.8$ | $1.7 \pm 0.9$ | $-86 \pm 8$ | $< 0.001$ | 3.93 |
| | EPSP 1 CV | $0.15 \pm 0.04$ | $0.53 \pm 0.21$ | $0.43 \pm 0.19$ | $+245 \pm 107$ | 0.001 | 1.94 |
| | EPSP 2 amp. | $8.9 \pm 4.8$ | $1.8 \pm 1.6$ | $2.7 \pm 1.9$ | $-85 \pm 11$ | $< 0.001$ | 3.32 |
| | PPR | $1.35 \pm 0.34$ | $1.60 \pm 0.79$ | $1.53 \pm 0.49$ | $+7 \pm 50$ | 0.502 | 0.24 |

**Figure 9.**
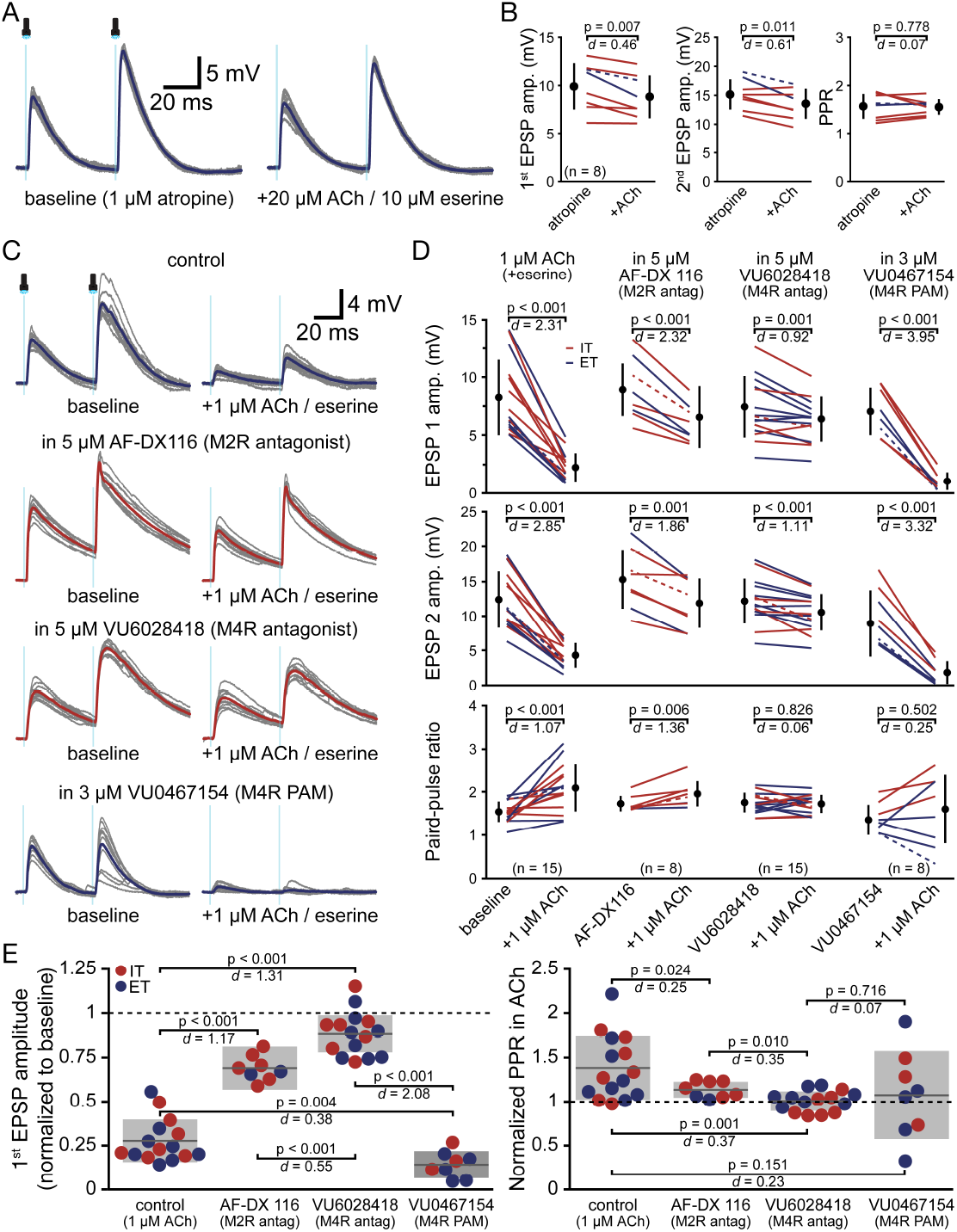
M4 muscarinic receptors mediate cholinergic suppression of commissural (COM) synaptic transmission. ***A***, Ten consecutive responses to optical activation of COM afferents (two 1-ms flashes at 20 Hz; grey traces) and averaged responses (blue traces) in an ET neuron in the presence of 20 µM atropine (a non-specific muscarinic receptor blocker; left) and after addition of 20 µM ACh (with 10 µM eserine; right). ***B***, Summary plots of EPSP amplitudes (left and middle) and paired-pulse ratios (PPR; right) in baseline conditions with atropine present and after addition of ACh for eight neurons. Individual neurons shown with lines, with means and standard deviations for the different conditions offset. Dashed lines indicate data from neuron shown in ***A. C***, Ten consecutive COM responses (grey) and averaged responses (colored traces) in IT (red) or ET (blue) neurons in various baseline conditions (as indicated; left) and after addition of 1 µM ACh (with 10 µM eserine; right). The top trace shows control experiments conducted in regular aCSF, while the other neurons were pre-exposed to M2- or M4-specific antagonists (middle traces) or an M4-specific positive allosteric modulator (PAM; lower traces), as indicated. ***D***, Plots of the effects of 1 µM ACh with eserine on COM EPSP amplitudes (top and middle) and paired-pulse ratios (PPR; bottom) for the different experimental conditions. Dashed lines indicate results from the neurons shown in ***C. E***, Plots of the normalized first EPSP amplitudes (relative to baseline EPSPs; left) and PPR (right) in the presence of ACh for the different experimental conditions. Grey lines show means, grey shaded areas indicate standard deviations. P-values (Student’s t tests) and effect sizes (Cohen’s *d*) for cross-group comparisons are shown.

Among the five mAChR subtypes, Gi/o-coupled M2 and M4 mAChRs are most associated with presynaptic regulation of transmitter release. To evaluate their potential involvement in regulating COM afferents, we tested the impact of 1 µM ACh (with 10 µM eserine) on COM EPSPs in the presence of M2- or M4-preferring pharmacological agents (**Figure 9C, D**). We chose to test a 1 µM concentration because it delivers substantial, but not saturating, inhibition of COM EPSPs (see **Figure 7**) that allows for observation of bidirectional changes in cholinergic modulation of COM transmission. It also allowed us to compare the impact of pharmacological manipulations with results from 15 experiments in which 1 µM ACh was applied first in a series of ACh concentrations (73 ± 12% and 65 ± 10% reductions of EPSPs 1 and 2, respectively, with an increase in PPR of 38 ± 36%; see **Figure 7**, above).

To preferentially block M2 receptors, we applied AF-DX 116 (5 µM; n = 8) for seven minutes prior to addition of 1 µM ACh. In the presence of the M2 antagonist, ACh reduced the first and second EPSPs by 31 ± 7 and 22 ± 10%, respectively, and increased PPR by 13 ± 9% (see **Figure 9D** for statistics). These changes were substantially less than those observed in neurons exposed to 1 µM ACh in the absence of antagonist (**Figure 9E**). However, preferential blockade of M4 receptors with VU6028418 (5 µM; n = 15) limited the effects of ACh to a greater extent, with EPSPs being reduced by only ~12% with no net changes in PPRs (0 ± 12%; **Figure 9D**; **Table 8**). Thus, in the presence of the M4 antagonist, ACh effects were far less than those observed with ACh alone or in the presence of the M2 antagonist (**Figure 9E**), and were similar to those observed in the presence of 1 µM atropine (see **Figure 9A**; p = 0.701, *d* = 0.15, when comparing ACh-induced changes in first EPSP amplitude in atropine and in VU6028418). Finally, we found that the M4-selective positive allosteric modulator (PAM), VU0467154 (3 µM; n = 8), enhanced the suppressive effect of 1 µM ACh on COM EPSPs, with first and second EPSPs being reduced by 86 ± 8% (p = 0.004, *d* = 0.38 when compared to 1 µM ACh alone) and 85 ± 11% (p < 0.001, *d* = 0.60), respectively (**Figure 9D, E**). In the presence of the M4 PAM, ACh-induced changes in PPR were highly variable (+7 ± 50%), likely because some EPSPs were reduced to almost undetectable levels, leading to large and variable differences in first and second EPSP measurements trial-to-trial and across neurons. Taken together, these data point to M4 receptors as being key mediators of cholinergic suppression of COM afferents in the PL cortex.

### Pharmacology of serotonergic modulation of COM afferents

Serotonergic inhibition of glutamatergic transmission in the forebrain is most associated with presynaptic 5-HT1B-(1B) type serotonin receptors (Tanaka and North, 1993; Rhoades et al., 1994; Laurent et al., 2002; Kjaerby et al., 2016; Rama, 2023), although 1A and 2A receptors have also been implicated (Troca-Marin and Geijo-Barrientos, 2010; Tian et al., 2017; Agahari and Stricker, 2021, 2025). The inclusion of 1A and 2A antagonists in our initial experiments (see **Figure 4**; **Table 6**) suggest that these receptors, that are otherwise prominent in the PL cortex (e.g., Amargos-Bosch et al., 2004), are not necessary for presynaptic regulation of glutamate release at COM terminals. Indeed, suppression of COM EPSPs by 40 µM 5-HT was similar in the presence or absence of blockers for 1A and 2A receptors (compare **Figures 4E, F** and **8B**). Because neuromodulation of glutamate release is most associated with signaling from Gi/o-coupled metabotropic receptors, we focused on Gi/o-coupled 5-HT1 and 5-HT5A (5A) receptor subtypes.

In control experiments in the absence of any serotonergic agents (n = 10), 2 µM 5-HT reduced the first and second EPSPs by 63 ± 10% and 51 ± 13%, respectively, increased PPRs by 34 ± 30%, and greatly (but with high variability) increased first EPSP CVs by over 100% (**Figure 10A, D**; **Table 9**). These effects were similar to those observed in our experiments in which neurons experienced 2 µM 5-HT as part of a series of 5-HT concentrations (see **Figure 8B**), where the first and second EPSPs were reduced by 63 ± 25% (n = 28; p = 0.931, *d* = 0.03, when comparing the 18 experiments in **Figure 8** to the 10 control experiments in **Figure 10**) and 61 ± 19% (p = 0.094, *d* = 0.60), respectively, and PPRs were increased by 16 ± 33% (p = 0.153, *d* = 0.57).

**Table 9.**
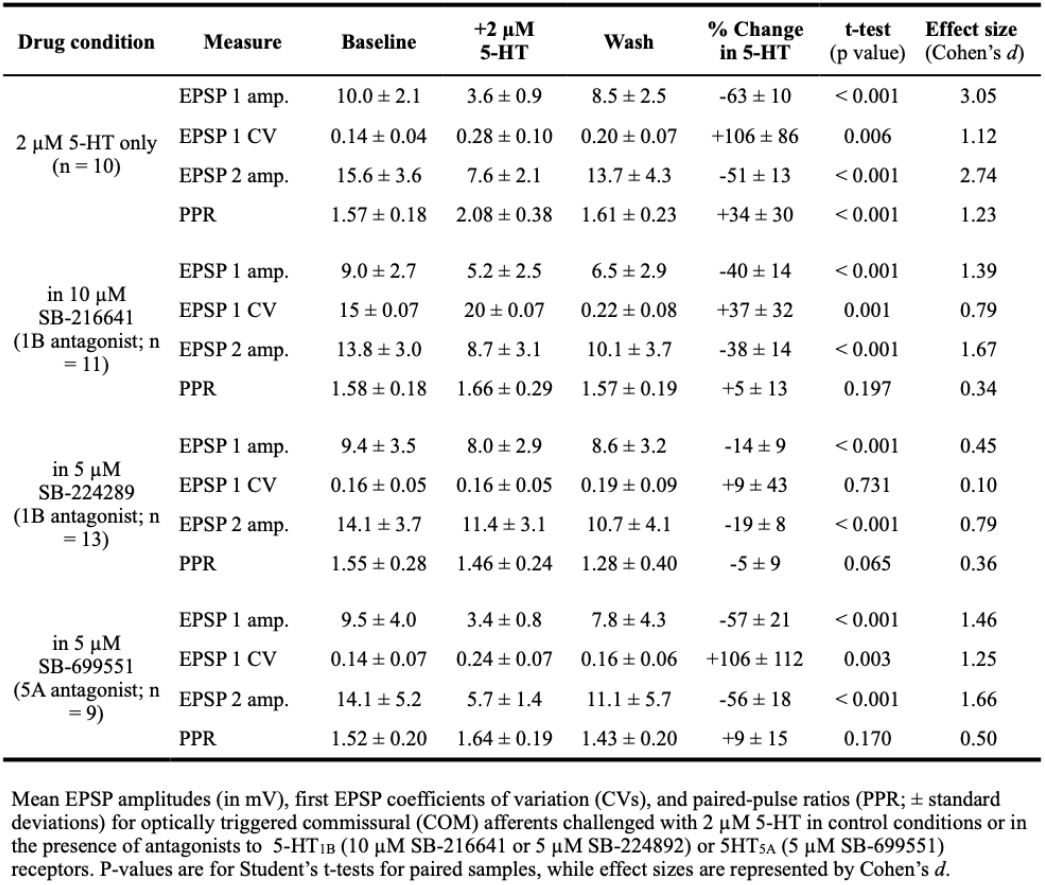
Impact of 5-HT1B blockade on serotonergic suppression of COM EPSPs.

| Drug condition | Measure | Baseline | +2 μM 5-HT | Wash | % Change in 5-HT | t-test (p value) | Effect size (Cohen’s *d*) |
| --- | --- | --- | --- | --- | --- | --- | --- |
| 2 μM 5-HT only ($n = 10$) | EPSP 1 amp. | $10.0 \pm 2.1$ | $3.6 \pm 0.9$ | $8.5 \pm 2.5$ | $-63 \pm 10$ | $< 0.001$ | 3.05 |
| | EPSP 1 CV | $0.14 \pm 0.04$ | $0.28 \pm 0.10$ | $0.20 \pm 0.07$ | $+106 \pm 86$ | 0.006 | 1.12 |
| | EPSP 2 amp. | $15.6 \pm 3.6$ | $7.6 \pm 2.1$ | $13.7 \pm 4.3$ | $-51 \pm 13$ | $< 0.001$ | 2.74 |
| | PPR | $1.57 \pm 0.18$ | $2.08 \pm 0.38$ | $1.61 \pm 0.23$ | $+34 \pm 30$ | $< 0.001$ | 1.23 |
| in 10 μM SB-216641 (1B antagonist; $n = 11$) | EPSP 1 amp. | $9.0 \pm 2.7$ | $5.2 \pm 2.5$ | $6.5 \pm 2.9$ | $-40 \pm 14$ | $< 0.001$ | 1.39 |
| | EPSP 1 CV | $15 \pm 0.07$ | $20 \pm 0.07$ | $0.22 \pm 0.08$ | $+37 \pm 32$ | 0.001 | 0.79 |
| | EPSP 2 amp. | $13.8 \pm 3.0$ | $8.7 \pm 3.1$ | $10.1 \pm 3.7$ | $-38 \pm 14$ | $< 0.001$ | 1.67 |
| | PPR | $1.58 \pm 0.18$ | $1.66 \pm 0.29$ | $1.57 \pm 0.19$ | $+5 \pm 13$ | 0.197 | 0.34 |
| in 5 μM SB-224289 (1B antagonist; $n = 13$) | EPSP 1 amp. | $9.4 \pm 3.5$ | $8.0 \pm 2.9$ | $8.6 \pm 3.2$ | $-14 \pm 9$ | $< 0.001$ | 0.45 |
| | EPSP 1 CV | $0.16 \pm 0.05$ | $0.16 \pm 0.05$ | $0.19 \pm 0.09$ | $+9 \pm 43$ | 0.731 | 0.10 |
| | EPSP 2 amp. | $14.1 \pm 3.7$ | $11.4 \pm 3.1$ | $10.7 \pm 4.1$ | $-19 \pm 8$ | $< 0.001$ | 0.79 |
| | PPR | $1.55 \pm 0.28$ | $1.46 \pm 0.24$ | $1.28 \pm 0.40$ | $-5 \pm 9$ | 0.065 | 0.36 |
| in 5 μM SB-699551 (5A antagonist; $n = 9$) | EPSP 1 amp. | $9.5 \pm 4.0$ | $3.4 \pm 0.8$ | $7.8 \pm 4.3$ | $-57 \pm 21$ | $< 0.001$ | 1.46 |
| | EPSP 1 CV | $0.14 \pm 0.07$ | $0.24 \pm 0.07$ | $0.16 \pm 0.06$ | $+106 \pm 112$ | 0.003 | 1.25 |
| | EPSP 2 amp. | $14.1 \pm 5.2$ | $5.7 \pm 1.4$ | $11.1 \pm 5.7$ | $-56 \pm 18$ | $< 0.001$ | 1.66 |
| | PPR | $1.52 \pm 0.20$ | $1.64 \pm 0.19$ | $1.43 \pm 0.20$ | $+9 \pm 15$ | 0.170 | 0.50 |

Mean EPSP amplitudes (in mV), first EPSP coefficients of variation (CVs), and paired-pulse ratios (PPR; ± standard deviations) for optically triggered commissural (COM) afferents challenged with 2 μM 5-HT in control conditions or in the presence of antagonists to 5-HT<sub>1B</sub> (10 μM SB-216641 or 5 μM SB-224892) or 5-HT<sub>5A</sub> (5 μM SB-699551) receptors. P-values are for Student’s *t*-tests for paired samples, while effect sizes are represented by Cohen’s *d*.

**Figure 10.**
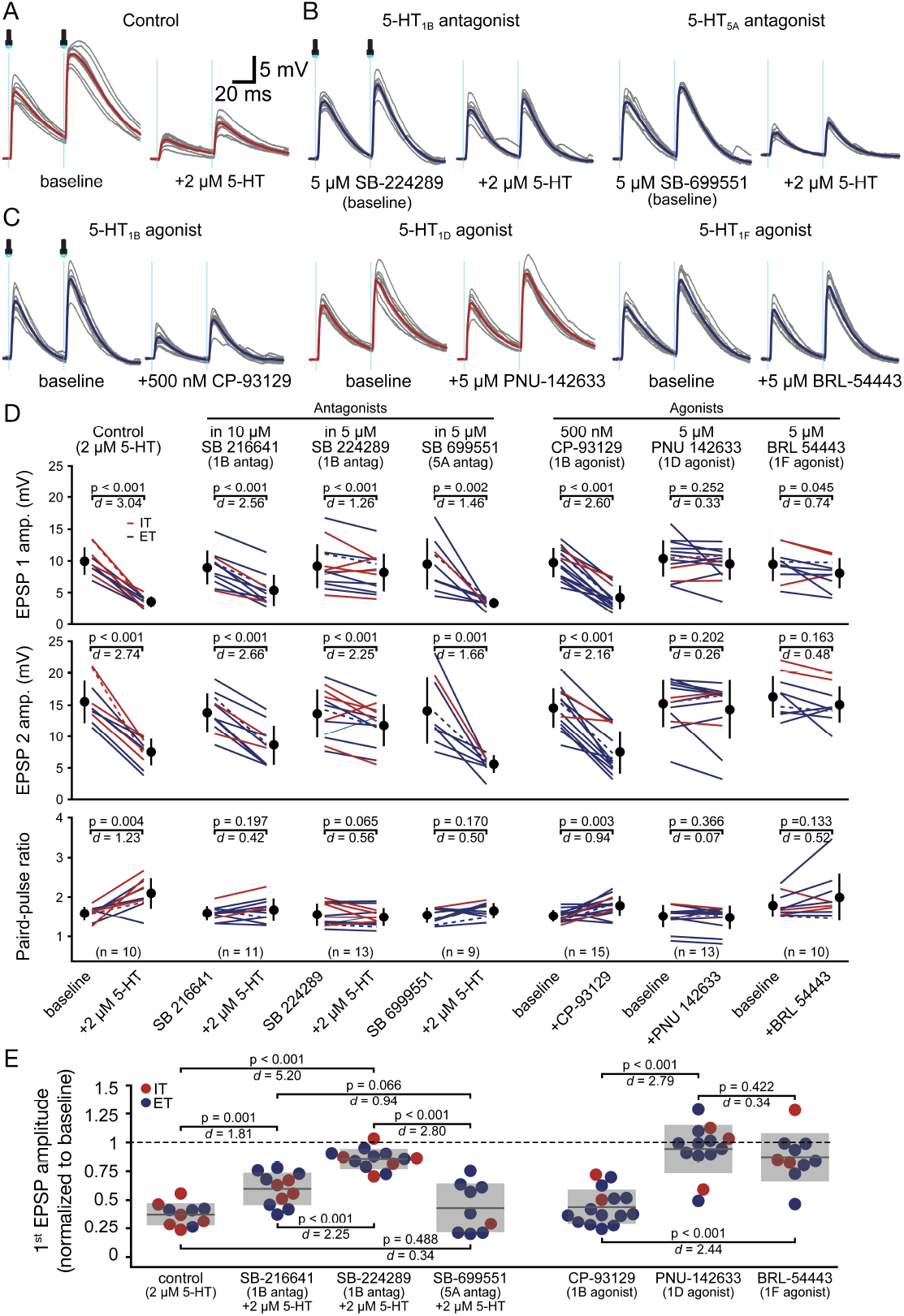
5-HT_1B_ receptors mediate sero-tonergic suppression of commissural (COM) synaptic transmission. ***A***, Ten consecutive responses to optical activation of COM afferents (two 1-ms flashes at 20 Hz; grey traces) and averaged responses (red traces) in an IT neuron in baseline conditions (left) and after addition of 2 µM 5-HT (right). Scale bar applies also to ***B*** and ***C. B***, Ten consecutive COM synaptic responses (grey) and averaged responses (blue traces) in ET neurons in antagonists for 5-HT_1B_ (left) or 5-HT_5A_ (right) receptors before and after the addition of 2 µM 5-HT. ***C***, Ten consecutive COM synaptic responses (grey) and averaged responses (colored traces) in ET (blue) and IT (red) neurons in baseline conditions and after addition of the indicated serotonergic agonists. ***D***, Plots of mean EPSP amplitudes (top and middle) and paired-pulse ratios (PPR; bottom) in baseline conditions and after the addition of 2 µM 5-HT (alone or in the presence of antagonists) or application of the indicated serotonergic agonists. Dashed lines indicate results from the neurons shown in ***A***-***C. E***, Plot of normalized first EPSP amplitudes (relative to baseline EPSPs) after the indicated pharmacological treatments. Grey lines show means, grey shaded areas indicate standard deviations. P-values (Student’s t tests) and effect sizes (Cohen’s *d*) for cross-group comparisons are shown.

The presence of 10 µM SB-216641 (n = 11), a 1B antagonist, modestly reduced the effect of 2 µM 5-HT on COM EPSPs, where the first and second EPSPs were reduced by 40 ± 14% and 38 ± 14%, respectively, concurrent with a small increase in PPR (+5 ± 13%; **Table 9**). However, 2 µM 5-HT was much less effective in suppressing COM EPSPs in the presence of a different 1B-preferring antagonist (SB-224289, 5 µM; n = 11), with reductions of the first and second EPSPs being < 20% and PPR being reduced rather than increased (**Figure 10B-E**; **Table 9**). Finally, a 5A receptor antagonist (SB-699551, 5 µM; n = 9) had negligible impact on serotonergic suppression of COM EPSPs (**Figure 10B, D, E**; **Table 9**).

When the effect of 2 µM 5-HT was compared across conditions, it’s efficacy in suppressing COM synaptic transmission was reduced by both 1B antagonists (most fully by SB-224289), but not by the 5A antagonist (**Figure 10E**). These data suggest 1B receptors may mediate serotonergic suppression of COM afferents. To test this further, in additional experiments we challenged COM EPSPs with agonists for 1B (CP-93129, 500 nM; n = 15), 5-HT1D (1D; PNU-102633, 5 µM; n = 13), or 5-HT1F (1F; BRL-54443, 5 µM; n= 10) receptors (**Figure 10C - E**; **Table 10**). Despite the applied concentration being an order of magnitude below that used for the 1D and 1F agonists, only the 1B agonist suppressed COM transmission to an extent comparable to 2 µM 5-HT, with initial EPSPs being reduced by 63 ± 10% and 56 ± 15% by 5-HT and CP-93129, respectively; n = 25; p = 0.209, *d* = 0.49, when compared to 2 µM 5-HT alone; see **Table 10**). Further, CP-93129 increased PPRs and the CV of initial EPSP amplitudes (**Figure 10D**; **Table 10**). These results further point to 1B receptors as being key regulators of glutamate release at COM terminals.

**Table 10.** Impact of 5-HT receptor agonists on COM EPSPs.

| Drug condition | Measure | Baseline | +agonist | Wash | % Change in agonist | t-test (p value) | Effect size (Cohen's $d$) |
| --- | --- | --- | --- | --- | --- | --- | --- |
| 500 nM CP-93129 (1B agonist; $n = 15$) | EPSP 1 amp. | $9.8 \pm 2.3$ | $4.3 \pm 1.8$ | $8.0 \pm 2.5$ | $-56 \pm 15$ | $< 0.001$ | 2.60 |
| | EPSP 1 CV | $0.16 \pm 0.04$ | $0.25 \pm 0.07$ | $0.16 \pm 0.04$ | $+62 \pm 62$ | 0.001 | 1.02 |
| | EPSP 2 amp. | $14.6 \pm 3.1$ | $7.6 \pm 3.4$ | $12.0 \pm 4.0$ | $-48 \pm 20$ | $< 0.001$ | 2.16 |
| | PPR | $1.50 \pm 0.15$ | $1.77 \pm 0.25$ | $1.50 \pm 0.20$ | $+18 \pm 19$ | 0.003 | 0.94 |
| 5 μM PNU-102633 (1D agonist; $n = 13$) | EPSP 1 amp. | $10.4 \pm 2.9$ | $9.6 \pm 2.5$ | $9.6 \pm 2.6$ | $-6 \pm 21$ | 0.252 | 0.33 |
| | EPSP 1 CV | $0.15 \pm 0.05$ | $0.15 \pm 0.04$ | $0.14 \pm 0.06$ | $0 \pm 0$ | 0.647 | 0.13 |
| | EPSP 2 amp. | $15.3 \pm 3.7$ | $14.3 \pm 4.6$ | $13.8 \pm 4.1$ | $-8 \pm 21$ | 0.202 | 0.26 |
| | PPR | $1.50 \pm 0.29$ | $1.47 \pm 0.30$ | $1.46 \pm 0.34$ | $-2 \pm 8$ | 0.366 | 0.07 |
| 5 μM BRL-54443 (1F agonist; $n = 10$) | EPSP 1 amp. | $9.5 \pm 2.6$ | $8.1 \pm 2.4$ | * | $-13 \pm 21$ | 0.045 | 0.56 |
| | EPSP 1 CV | $0.13 \pm 0.05$ | $0.14 \pm 0.04$ | * | $+10 \pm 29$ | 0.568 | 0.15 |
| | EPSP 2 amp. | $16.3 \pm 3.3$ | $15.1 \pm 2.9$ | * | $-6 \pm 16$ | 0.163 | 0.40 |
| | PPR | $1.77 \pm 0.29$ | $1.98 \pm 0.59$ | * | $+11 \pm 18$ | 0.133 | 0.44 |

Mean EPSP amplitudes (in mV), first EPSP coefficients of variation (CVs), and paired-pulse ratios (PPR; ± standard deviations) for optically triggered commissural (COM) afferents challenged with agonists for 5-HT<sub>1B</sub> (CP-93129; 500 nM), 5-HT<sub>1D</sub> (PNU-102633; 5 μM), or 5-HT<sub>1F</sub> (BRL-54443; 5 μM) receptors. \*For BRL-54443, there was no apparent effect of the agonist and so additional manipulations were done instead of a wash. P-values are for Student's t-tests for paired samples, while effect sizes are represented by Cohen's $d$ .

### Modulation of COM afferents with presynaptic DREADD receptors

To directly assess the ability of presynaptic metabotropic receptors to modulate glutamate release from COM afferents, we expressed Gi/o-(hM4Di) or Gq-(hM3Dq) coupled DREADDs (“designer receptors exclusively activated by designer drugs”) with ChR2 unilaterally in the PL cortex (**Figure 11A, D**) and measured optically evoked EPSPs in contralateral IT and ET target neurons in baseline conditions and after adding the DREADD-specific agonist CNO (50 nM, 500 nM, or 5 µM).

**Figure 11.**
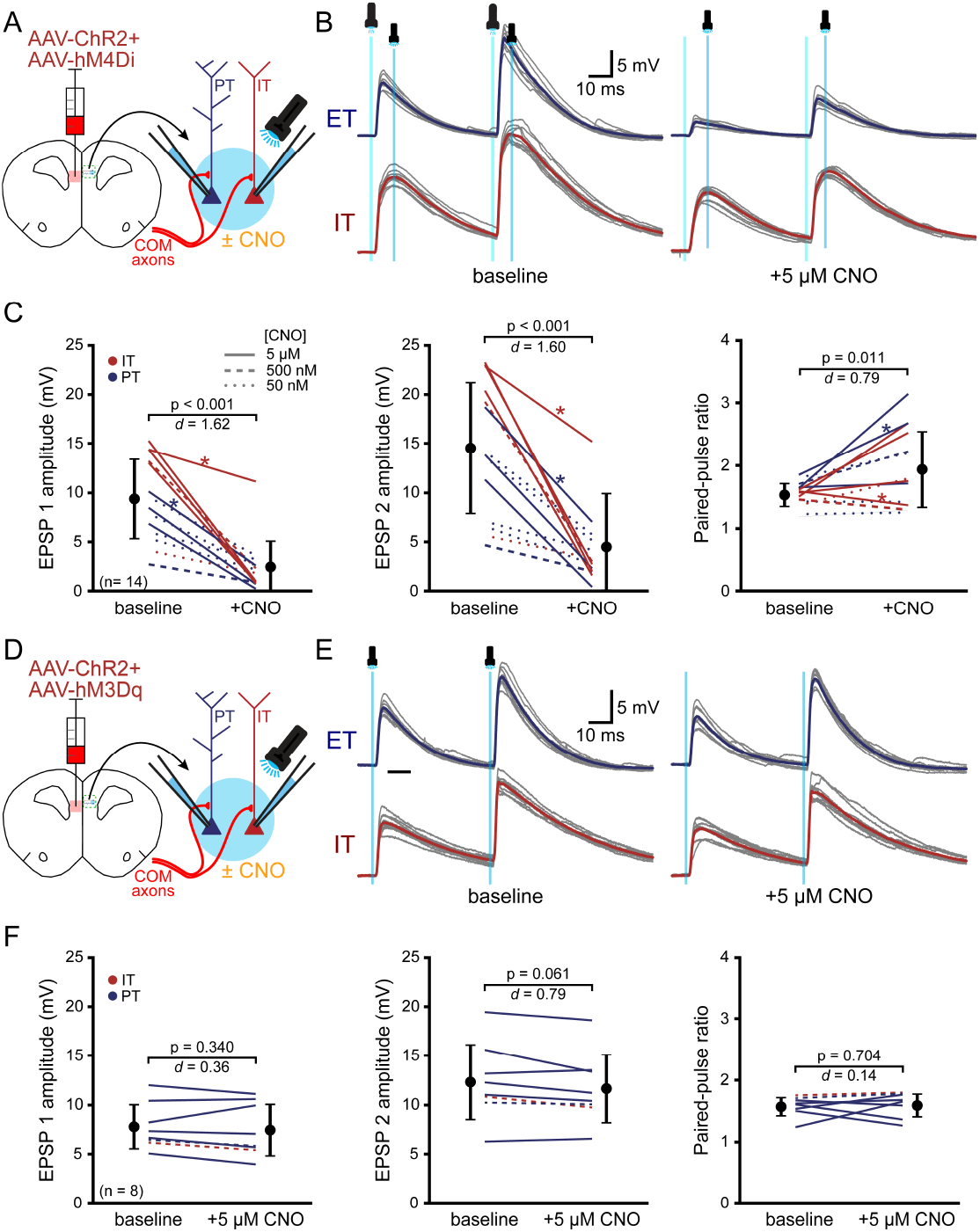
Activation of G_i/o_, but not G_q_, presynaptic metabotropic receptors suppresses commissural (COM) synaptic transmission. ***A***, Diagram of experimental setup, with AAV-induced co-expression of channel-rhodopsin-2 (ChR2) and hM4Di (G_i/o_-coupled) receptors in one cerebral hemisphere and subsequent optical activation of COM afferents in the opposite hemisphere before and after the application of clozapine-N-oxide (CNO). ***B***, Ten consecutive responses to pairs (20 Hz) of COM afferent activations (grey traces) and their averaged responses (colored traces) in a pair of IT (red) and ET (blue) neurons in baseline conditions (left) and after addition of 5 µM CNO (right). ***C***, Comparisons of EPSP amplitudes (left and middle) and paired-pulse ratios (PPR; right) before and after CNO applications at the noted concentrations for 14 neurons. Mean responses (± standard deviations) shown with offset symbols. P-values (Student’s t-tests for paired data) and effect sizes (Cohen’s *d*) shown for each. Asterisks indicate data from panel ***B. D***-***F***, Similar to ***A***-***C***, but for co-expression of ChR2 and hM3Dq, a G_q_-coupled receptor for CNO. In panel ***F***, dashed lines indicate data from experiments shown in panel ***E***.

With hM4Di co-expressed with ChR2, application of 5 µM CNO frequently resulted in profound and sometimes near-complete inhibition of optically evoked EPSPs (**Figure 11C**) that did not wash out after many minutes. Still, as observed in simultaneous recordings of COM EPSPs in an IT-ET neuron pair (**Figure 11B**), results with 5 µM CNO could be variable even under identical experimental conditions. This likely reflects stochastic differences in hM4Di expression levels across ChR2-expressing presynaptic COM terminals. Because 5 µM CNO generated long-lasting, and essentially irreversible effects, in some experiments we lowered the concentration of CNO to 500 or 50 nM to see if we could reverse CNO-induced effects during a 15-minute wash (**Figure 11C**). However, across all CNO concentrations tested, effects on COM EPSPs were long-lasting and failed to reverse. When data for all CNO concentrations were combined (n = 14), CNO suppressed the first and second EPSPs by 72 ± 22% and

68 ± 21%, respectively, and increased PPR by 26 ± 32% (**Table 11**). These data confirm that presynaptic Gi/o-coupled receptors can be powerful modulators of synaptic transmission at COM afferents.

**Table 11.** Presynaptic chemogenetic modulation of COM afferents.

| DREADD receptor | Measurement | Baseline | +CNO<br>(0.05 - 5 μM) | Change in<br>CNO (%) | CNO effect | |
| --- | --- | --- | --- | --- | --- | --- |
|  |  |  |  |  | t-test<br>(p value) | Effect size<br>(Cohen's *d*) |
| hM4Di<br>(n = 14) | EPSP 1 amp. | 9.4 ± 4.1 | 2.5 ± 2.7 | -72 ± 22% | < 0.001 | 1.62 |
| | EPSP 1 CV | 0.17 ± 0.07 | 0.57 ± 0.29 | +253 ± 202 | < 0.001 | 1.54 |
| | EPSP 2 amp | 14.6 ± 6.8 | 4.1 ± 3.6 | -68 ± 21 | < 0.001 | 1.60 |
| | PPR | 1.54 ± 0.20 | 1.95 ± 0.62 | +26 ± 32 | 0.011 | 0.80 |
| hM3Dq<br>(n = 8) | EPSP 1 amp. | 7.8 ± 2.3 | 7.4 ± 2.7 | -6 ± 13 | 0.340 | 0.13 |
| | EPSP 1 CV | 0.17 ± 0.06 | 0.17 ± 0.04 | +5 ± 18 | 0.789 | 0.07 |
| | EPSP 2 amp | 12.4 ± 3.9 | 11.7 ± 3.6 | -5 ± 6 | 0.061 | 0.18 |
| | PPR | 1.58 ± 0.16 | 1.61 ± 0.20 | +3 ± 16 | 0.704 | 0.14 |

Mean EPSP amplitudes (in mV), first EPSP coefficients of variation (CVs), and paired-pulse ratios (PPR; ± standard deviations) for optically triggered commissural (COM) afferents expressing presynaptic hM4Di or hM3Dq receptors in baseline conditions and after exposure to clozapine-N-oxide (CNO). Effects of CNO did not wash out within 15 minutes. P-values are for Student's t-tests for paired samples. Effect sizes are represented by Cohen's *d*.

Finally, we tested whether activation of presynaptic Gq receptors might modulate COM synaptic transmission by unilaterally co-expressing hM3Dq with ChR2 in PL cortex and measuring optically evoked COM EPSPs in baseline conditions and after the addition of CNO (**Figure 11D, E**). Unlike in the hM4Di experiments described above, CNO application at 5 µM had negligible impact on COM transmission (n = 8; **Figure 11E, F**; **Table 11**), with EPSP amplitudes (mean changes of −6 + 13% and −5 ± 6% for the first and second EPSPs, respectively) and PPRs (+3 ± 16%) being very similar to baseline values (**Table 11**). These data suggest that presynaptic expression of Gq-coupled receptors may have negligible impact on synaptic release at COM terminals. However, a limitation of our approach is that we cannot confirm that hM3Dq traffics to COM axon terminals, as both hM3Dq and ChR2 in our experiments were tagged with the same fluorophore (mCherry). Whereas a positive result with CNO would be readily interpretable (as in the hM4Di experiment above), the lack of efficacy of CNO in our hM3Dq experiment does not preclude potential presynaptic Gq-mediated modulation of synaptic transmission, as was recently shown for Gq-coupled 2A receptors at thalamocortical terminals in the retrosplenial cortex (Ekins et al., 2025).

However, this experiment serves as an effective control by demonstrating that CNO does not have non-specific effects that impact optically evoked COM EPSPs.

## Discussion

Previous studies have found that 5-HT and ACh differentially regulate the postsynaptic excitability of IT and ET neurons in the mouse cortex (Dembrow et al., 2010; Avesar and Gulledge, 2012; Stephens et al., 2014; Joshi et al., 2015; Baker et al., 2018), suggesting that these neuromodulators may shift the balance of cortical output toward cortical and striatal targets or deep brain structures, respectively. In the present study, we tested whether presynaptic regulation of excitatory synaptic transmission in the PL cortex is similarly afferent-and/ or target-specific. As discussed below, our results reveal target-independent selective presynaptic modulation of COM terminals, with both ACh and 5-HT preferentially gating (i.e., being permissive of) MDN inputs that selectively target IT neurons in layer 5 and drive local network activity to also excite ET neurons.

### Targeting of COM and MDN afferents

Consistent with several previous studies (Lee et al., 2014; Dembrow et al., 2015; Anastasiades et al., 2018; Leyrer-Jackson and Thomas, 2019; Yi et al., 2022), we found that optogenetic activation of COM afferents generated robust and roughly equivalent monosynaptic responses in IT and ET target neurons (see **Figures 1D** and **2B**). However, unlike prior studies, we found that COM synapses displayed largely target-independent short-term plasticity (paired-pulse facilitation) across a range of input frequencies (10 to 40 Hz; see **Figures 1E** and **3B**). This contrasts with earlier reports that COM afferents exhibit paired-pulse facilitation in ET neurons but paired-pulse depression in IT neurons (Lee et al., 2014; Leyrer-Jackson and Thomas, 2019), less paired-pulse depression in ET neurons (Yi et al., 2022), or paired-pulse depression of similar magnitudes at unitary corticocortical connections involving presynaptic IT neurons (Morishima et al., 2011). The diversity of these prior results cannot be easily explained by differences in animal age or strain (adult C57 mice, except for Morishima et al.) or by differences in extracellular calcium concentrations, which were 2 mM (Lee et al.; Yi et al.) or lower (~ 1 mM; Morishima et al.; Leyrer-Jackson and Thomas) and therefore expected to promote facilitating synaptic responses. Interestingly, all of these previous studies used 10 Hz stimulation, which we found to generate less facilitation than EPSPs driven at 20 or 40 Hz (see **Figure 3B** and Martinetti et al., 2022), and employed shorter inter-trial intervals (8 - 10 s) than used in our experiments (15 s). These differences in experimental approach may contribute to the diversity of prior findings, as Burke et al. (2018) observed in layer 5 of mPFC a paired-pulse facilitation of similar magnitude to our study (PPR of ~1.5) for a range of afferent input, including COM EPSPs and unitary events, with 20 Hz stimulation delivered at 15 s intervals. Our data go further than these earlier studies in showing that short-term synaptic plasticity of COM afferents in the adult mouse PL cortex is largely target-independent in layer 5 across the broader range of 10 to 40 Hz.

We found thalamocortical projections from the MDN to be highly selective for IT target neurons in the PL cortex. This is broadly consistent with the results of Collins et al. (2018), who observed a strong bias for monosynaptic connections in IT target neurons. However, unlike Collins et al., who reported paired-pulse depression of MDN inputs to layer 2/3 pyramidal neurons at 10 Hz, we observed facilitation in layer 5 IT neurons across a range of frequencies (see **Figure 3B**). This difference may reflect target-neuron-dependent differences in short-term plasticity (as MDN-driven PPR in layer 5 IT neurons has not been previously reported), but may also reflect subtle differences in experimental approach (see above).

### Afferent-selective neuromodulation of excitatory drive

Given that ACh and 5-HT reciprocally regulate the postsynaptic excitability of IT and ET neurons, we tested whether these neuromodulators might also differentially regulate excitatory synaptic drive in IT and ET neurons, perhaps in an afferent- and/or target-specific manner. Our main result is that both ACh and 5-HT suppressed monosynaptic glutamate release selectively in COM afferents, albeit to different degrees, while having little impact on monosynaptic MDN transmission.

Although afferent-specific cholinergic modulation of glutamate release has been reported in the PL cortex (Banks et al., 2021), sensory cortex (Gil et al., 1997; Hsieh et al., 2000), hippocampus (Kahle and Cotman, 1989; Hasselmo and Schnell, 1994; Kremin and Hasselmo, 2007), olfactory tubercle (Owen and Halliwell, 2001), piriform cortex (Hasselmo and Bower, 1992; Linster et al., 1999), and amygdala (Tryon et al., 2023), no prior study has demonstrated presynaptic cholinergic regulation of an optogenetically isolated afferent in the neocortex. Our results, which also include first-ever tests of cholinergic or serotonergic modulation of monosynaptic connections isolated with TTX and 4-AP, are consistent with the hypotheses that ACh acts presynaptically to suppress corticocortical network activity while being permissive for long-range excitatory afferents (for review, see Giocomo and Hasselmo, 2007).

Afferent-specific serotonergic suppression of transmission has been described previously for COM, relative to MDN, inputs to layer 5 neurons in PL cortex, albeit without identifying target neuron subtypes (Kjaerby et al., 2016). Conversely, in the nucleus accumbens, 5-HT was reported to selectively suppress thalamic inputs but not afferents from the mPFC (Christoffel et al., 2021). This suggests that presynaptic 5-HT receptor expression may be limited to a subpopulation of bilaterally projecting IT neurons in the mPFC.

We found cholinergic suppression of COM transmission in the PL cortex to be almost absolute (>85%), suggesting that the majority of COM terminals express M4 receptors. Serotonergic modulation of COM EPSPs was less robust (~55% reduction at 40 µM 5-HT; see also Kjaerby et al., 2016) and more variable (SDs of 18% and 6%, for 5-HT and ACh, respectively). More modest effects of 5-HT (~25% reduction) were reported for electrically evoked (but putative COM) synaptic transmission in pyramidal neurons in the anterior cingulate cortex (Troca-Marin and Geijo-Barrientos, 2010; see also Tian et al., 2017).

Strong cholinergic, but moderate serotonergic suppression of glutamatergic transmission may be a conserved feature of presynaptic neuromodulation across brain regions. For instance, in the olfactory tubercle both ACh and 5-HT exhibit afferent-specific presynaptic regulation of glutamate release at local associational synapses, but the suppressive effect of ACh (~70% suppression with an ~60% increase in PPR) is much greater than that for 5-HT (~30% suppression with an ~20% increase in PPR; Owen and Halliwell, 2001; Hadley and Halliwell, 2010). Thus, there may be distinct presynaptic mechanisms that allow ACh and 5-HT to regulate synaptic transmission to different degrees. Alternatively, and consistent with the results of Christoffel et al. (2021), it is possible that there is heterogeneity among the IT neurons that contribute to COM projections, with only a subset of neurons expressing presynaptic 1B receptors.

### Neuromodulation of COM afferents is presynaptic

Our results are consistent with the hypothesis that ACh and 5-HT act presynaptically to limit glutamate release at COM terminals. First, although it is expected that postsynaptic changes in membrane excitability will impact postsynaptic potentials (e.g., by changing the neuron’s electrotonic structure), such changes should have broadly similar effects on both COM and MDN EPSPs, as both afferents innervate similar dendritic domains (Anastasiades and Carter, 2021). Second, the M4 receptors responsible for cholinergic suppression of COM transmission are not associated with postsynaptic regulation of pyramidal neurons (Gulledge et al., 2009; Dasari and Gulledge, 2011), whereas 5-HT reduced COM EPSPs approximately equally in the presence or absence of antagonists for the 1A and 2A receptors responsible for the opposing postsynaptic serotonergic effects in ET and IT neurons. Third, while 20 µM ACh reduced COM EPSPs in ET neurons by almost 90% (see **Table 5**), it also increases spontaneous (i.e., non-evoked) EPSP amplitude and frequency in ET neurons under the same experimental conditions (Gulledge, 2024). Finally, ACh across a range of concentrations increased the CV of EPSP amplitudes and promoted paired-pulse facilitation, a classic, but not definitive (see Burke et al., 2018), indicator of presynaptic modulation of transmitter release.

### Functional implications

Although both 5-HT and ACh suppress excitatory transmission at COM terminals, the magnitude of suppression is greater for ACh (**Table 5**) than for 5-HT (**Table 6**). Combined with target-specific postsynaptic regulation of IT and ET excitability, these data suggest that 5-HT and ACh may differentially impact information processing in cortical circuits. **Figure 12** presents a model for serotonergic and cholinergic regulation of the PL cortex that incorporates both pre- and postsynaptic mechanisms in a subset of local and long-distance circuits. In **Figure 12A**, IT neurons innervate both IT and ET target neurons, while ET neurons selectively target other ET neurons (Morishima and Kawaguchi, 2006; Morishima et al., 2011; Kiritani et al., 2012; Gulledge, 2024; but see Bodor et al., 2025). IT neurons also project bilaterally to the striatum with dense terminal ramifications, while ET neurons project only unilaterally and with fewer, more focused axon collaterals to the striatum (Zheng and Wilson, 2002; Hooks et al., 2018). Recent connectivity mapping has found that the MDN and PL cortex project to similar striatal target zones in the dorsomedial stratum and nucleus accumbens (Hunnicutt et al., 2016), and that these striatal circuits, in turn, provide feedback to the MDN (for review, see Groenewegen et al., 2016).

**Figure 12.**
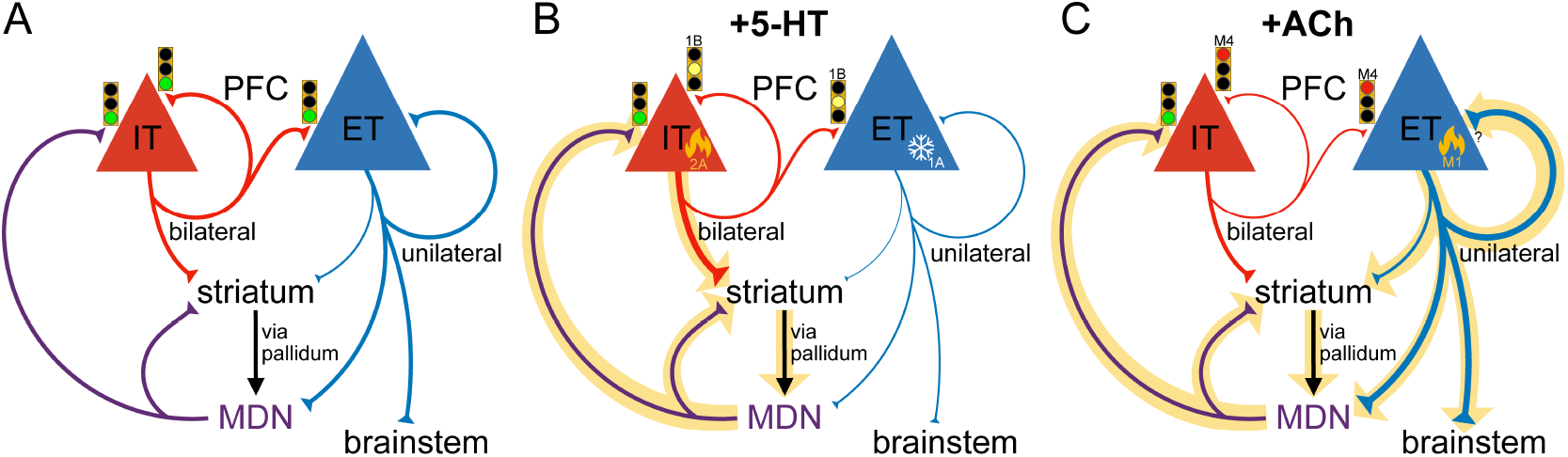
Circuit-level impact of 5-HT and ACh release in the prefrontal cortex. ***A***, Diagram of local and long-distance circuits subserved by IT (red) and ET (blue) neurons in the prefrontal cortex (PFC). IT neurons project bilaterally within the cortex and to the striatum (both the dorsomedial striatum and nucleus accumbens), whereas networks of interconnected ET neurons project ipsilaterally within the cortex, moderately to the striatum, and to one or more subcortical targets (e.g., the brainstem and/or the mediodorsal nucleus [MDN] of the thalamus). The MDN is bidirectionally connected with the striatum (projecting direct efferents and receiving feedback via the pallidum) and with the cortex (selectively targeting IT neurons in the PFC and receiving feedback via ET projections and striatal circuits). Green traffic lights indicate normal synaptic transmission in the absence of neuromodulation. ***B***, 5-HT increases the gain of IT neurons via 2A receptors and strongly inhibits ET neurons via 5-HT_1A_ (1A) receptors, while moderately suppressing IT-derived corticocortical communication (yellow traffic symbols), via presynaptic 5-HT_1B_ (1B) receptors. The net effect is preferential amplification of thalamocortically driven corticostriatal output bilaterally, which in turn may provide feedback to the MDN via the pallidum. ***C***, In the presence of ACh, M4 muscarinic acetylcholine receptors strongly suppress IT-derived corticocortical communication (red traffic lights) while M1 muscarinic receptors preferentially enhance the gain of interconnected ET neurons, promoting corticofugal output to the ipsilateral striatum and deep structures, including the MDN, which in turn promotes bilateral corticostriatal output via IT neurons.

In this model, cortical release of 5-HT is expected to increase the gain of IT neurons (via postsynaptic 2A receptors), silence ET neurons (via postsynaptic 1A receptors), and reduce, but not eliminate, IT-mediated corticocortical transmission via presynaptic 1B receptors (**Figure 12B**). Therefore, 5-HT release in the cortex may preferentially promote corticostriatal circuits driven primarily by thalamocortical MDN afferents that are not sensitive to presynaptic serotonergic regulation. The increased gain of IT neurons may compensate for the reduced corticocortical transmission to amplify, and perhaps widen, cortico-striatal-thalamic circuits.

Corticocortical transmission is more sensitive to cholinergic modulation (via presynaptic M4 receptors; **Figure 12C**), which also preferentially enhances the excitability of, and recurrent activity in, networks of ET neurons via postsynaptic M1 receptors (Gulledge et al., 2009; Baker et al., 2018; Gulledge, 2024). Given that a subpopulation of ET neurons monosynaptically excites thalamocortical neurons in the MDN (Collins et al., 2018), ACh release in the cortex may directly promote behavioral output (via brainstem projections) by driving focused, and perhaps narrowing, thalamocortical loops (**Figure 12C**) that have more balanced, and perhaps overlapping, contributions of ET and IT corticostriatal projections, but strongly reduced intra-areal corticocortical communication (Shepherd and Yamawaki, 2021). Although greatly oversimplified and omitting many neuron subtypes (including GABAergic interneurons) and synapses that may also be modulated by ACh and/ or 5-HT, the model presented in **Figure 12** provides testable predictions about corticostriatal circuit performance across a variety of neuromodulatory conditions that are relevant for understanding normal and pathological decision making and behavior.

